# Hormonal correlates of pathogen disgust: Testing the Compensatory Prophylaxis Hypothesis

**DOI:** 10.1101/156430

**Authors:** Benedict C Jones, Amanda C Hahn, Claire I Fisher, Hongyi Wang, Michal Kandrik, Anthony J Lee, Joshua Tybur, Lisa DeBruine

**Author notes:** **Corresponding author:** Benedict C Jones, Institute of Neuroscience & Psychology, University of Glasgow, UK.

## Abstract

Raised progesterone during the menstrual cycle is associated with suppressed physiological immune responses, reducing the probability that the immune system will compromise the blastocyst’s development. The Compensatory Prophylaxis Hypothesis proposes that this progesterone-linked immunosuppression triggers increased disgust responses to pathogen cues, compensating for the reduction in physiological immune responses by minimizing contact with pathogens. Although a popular and influential hypothesis, there is no direct, within-woman evidence for correlated changes in progesterone and pathogen disgust. To address this issue, we used a longitudinal design to test for correlated changes in salivary progesterone and pathogen disgust (measured using the pathogen disgust subscale of the Three Domain Disgust Scale) in a large sample of women (N=375). Our analyses showed no evidence that pathogen disgust tracked changes in progesterone, estradiol, testosterone, or cortisol. Thus, our results provide no support for the Compensatory Prophylaxis Hypothesis of variation in pathogen disgust.

## Introduction

Suppressed physiological immune responses during the luteal phase of the menstrual cycle (when raised progesterone prepares the body for pregnancy) reduce the probability that the immune system will compromise the development of the blastocyst (reviewed in Fleischman & Fessler, 2011). The Compensatory Prophylaxis Hypothesis proposes that this progesterone-linked immunosuppression is associated with increased disgust toward pathogen cues, compensating for the reduction in physiological immune responses by reducing the probability of contact with pathogens (Fessler et al., 2005; Fleischman & Fessler, 2011; Żelaźniewicz et al., 2016).

Fleischman and Fessler (2011) and Żelaźniewicz et al. (2016) have presented the strongest evidence for (and most direct tests of) the Compensatory Prophylaxis Hypothesis to date. In both studies, women with higher progesterone levels reported stronger disgust toward pathogen cues. Another study reporting stronger disgust responses to pathogen cues during the first (i.e., highest-progesterone) trimester of pregnancy has also been interpreted as supporting the Compensatory Prophylaxis Hypothesis (Fessler et al.,2005). However, these three studies employed between-subject designs, which have been shown to be weak (e.g., underpowered) tests for hypotheses concerning hormone-linked changes in behavior (Gangestad et al., 2016) and allow only indirect tests of the hypothesis that within-woman changes in pathogen disgust and progesterone are correlated. The two studies that measured progesterone levels (Fleischman & Fessler, 2011; Żelaźniewicz et al., 2016) employed relatively small sample sizes (Ns of 79 and 30, respectively), meaning that they were likely underpowered (see Gangestad et al., 2016).

Other studies often cited as evidence for the Compensatory Prophylaxis Hypothesis are also problematic. For example, greater hostility to out-group individuals during the first trimester of pregnancy has been interpreted as evidence for the Compensatory Prophylaxis Hypothesis because out-group individuals putatively pose a greater pathogen threat than do in-group individuals (Navarette et al., 2007). However, the hypothesis that hostility to out-group individuals necessarily reflects pathogen avoidance has recently been extensively critiqued (Aarøe et al., 2016; de Barra & Curtis, 2012; Tybur et al., 2016). Reports that women show stronger aversions to individuals displaying facial cues of illness (e.g., pallor) at high-progesterone points in the menstrual cycle (Jones et al., 2005) have also been interpreted as evidence for the Compensatory Prophylaxis Hypothesis. These results were not replicated in a higher-powered study that directly tested for correlated changes in measured progesterone and aversions to facial cues of illness (Jones et al., 2017a).

In summary, considering its influence in both the emotion and endocrinology literatures, the Compensatory Prophylaxis Hypothesis is supported by a surprisingly weak body of evidence. In the current study, we rigorously tested the Compensatory Prophylaxis Hypothesis by using a longitudinal design to investigate whether within-woman changes in steroid hormone levels (including progesterone) and changes in components of disgust sensitivity (including pathogen disgust) were correlated in a large sample of women (N=375). We assessed disgust sensitivity using Tybur et al.’s (2009) Three Domain Disgust Scale, which assesses disgust sensitivity in three different domains: pathogen disgust, sexual disgust, and moral disgust. The Compensatory Prophylaxis Hypothesis predicts that pathogen disgust will track changes in women’s progesterone levels (Fessler et al., 2005; Fleischman & Fessler, 2011; Żelaźniewicz et al., 2016). Indeed, the studies testing the Compensatory Prophylaxis Hypothesis have each used either the Three Domain Disgust Scale, similar self-report measures of disgust or contamination sensitivity, or disgust ratings of images portraying cues to pathogens (Fessler et al., 2005; Fleischman & Fessler, 2011; Żelaźniewicz et al., 2016).

## Methods

### Participants

We tested 375 heterosexual women (mean age=21.6 years, SD=3.3 years), all of whom reported that they were not using any form of hormonal contraceptive (i.e., reported having natural menstrual cycles). Participants completed up to three blocks of test sessions. Each of the three blocks of test sessions consisted of five weekly test sessions. Women participated as part of a large study of possible effects of steroid hormones on women’s behavior (Jones et al., 2017a). The data analyzed here are all responses from blocks of test sessions where women were not using any form of hormonal contraceptive and test sessions where they completed at least one subscale of Tybur et al.’s (2009) Three Domain Disgust Scale. Following these restrictions, 337 women had completed five or more test sessions and 98 of these women completed ten test sessions. Thirty-eight women completed fewer than five test sessions.

### Three Domain Disgust Scale

Participants completed Tybur et al.’s (2009) Three Domain Disgust Scale in each test session. This 21-item measure asks participants to rate each of 21 actions from not at all disgusting (0) to extremely disgusting (6). The actions were divided into three domains: pathogen disgust (e.g., stepping on dog poop), sexual disgust (e.g., hearing two strangers having sex), and moral disgust (e.g., deceiving a friend). Question order was fully randomized. The full instructions for the questionnaire were: “The following items describe a variety of concepts. Please rate how disgusting you find the concepts described in the items, where 0 means that you do not find the concept disgusting at all, and 6 means that you find the concept extremely disgusting.”

The mean score on the pathogen disgust subscale was 25.99 (SD=7.98), the mean score on the sexual disgust subscale was 19.95 (SD=8.71), and the mean score on the moral disgust subscale was 27.82 (SD=8.32). Intra-class correlation coefficients were high for each subscale (pathogen: .82; 95% CIs: .80, .85; sexual: .88; 95% CIs: .86, .89; moral=.79; 95% CIs: .76, .82). Consistent with past research (Olatunji et al., 2012), these intra-class correlation coefficients indicate that scores on the Three Domain Disgust Scale are stable over time (or, at least, over the time span sampled in the current study). Nevertheless, small fluctuations in disgust sensitivity could covary with variation in hormones.

### Saliva samples

Participants provided a saliva sample via passive drool (Papacosta & Nassis, 2011) in each test session. Participants were instructed to avoid consuming alcohol and coffee in the 12 hours prior to participation and avoid eating, smoking, drinking, chewing gum, or brushing their teeth in the 60 minutes prior to participation. Each woman’s test sessions took place at approximately the same time of day to minimize effects of diurnal changes in hormone levels (Veldhuis et al., 1988; Bao et al., 2003).

Saliva samples were frozen immediately and stored at −32°C until being shipped, on dry ice, to the Salimetrics Lab (Suffolk, UK) for analysis, where they were assayed using the Salivary 17β-Estradiol Enzyme Immunoassay Kit 1-3702 (M=3.30 pg/mL, SD=1.27 pg/mL, sensitivity=0.1 pg/mL, intra-assay CV=7.13%, inter-assay CV=7.45%), Salivary Progesterone Enzyme Immunoassay Kit 1-1502 (M=148.59 pg/mL, SD=96.20 pg/mL, sensitivity=5 pg/mL, intra-assay CV=6.20%, inter-assay CV=7.55%), Salivary Testosterone Enzyme Immunoassay Kit 1-2402 (M=87.57 pg/mL, SD=27.19 pg/mL, sensitivity<1.0 pg/mL, intra-assay CV=4.60%, inter-assay CV=9.83%), and Salivary Cortisol Enzyme Immunoassay Kit 1-3002 (M=0.23 μg/dL, SD=0.16 μg/dL, sensitivity<0.003 μg/dL, intra-assay CV=3.50%, inter-assay CV=5.08%). Although Fleischman and Fessler (2011) and Żelaźniewicz et al. (2016) only reported progesterone in their studies, we measured and report analyses of estradiol, cortisol, and testosterone, in addition to progesterone, to test whether links between pathogen disgust and hormonal status are driven specifically by progesterone, as the Compensatory Prophylaxis Hypothesis proposes. Mean minimum and maximum hormone levels are given in our Supplemental Information.

Hormone levels more than three standard deviations from the sample mean for that hormone or where Salimetrics indicated levels were outside the sensitivity range of their relevant ELISA were excluded from the dataset (~1 % of hormone measures were excluded for these reasons). The descriptive statistics given above do not include these excluded values. Values for each hormone were centered on their subject-specific means to isolate effects of within-woman changes in hormones. They were then scaled (i.e., divided by a constant) so the majority of the distribution for each hormone varied from -.5 to .5 to facilitate calculations in the linear mixed models. Since hormone levels were centered on their subject-specific means, women with only one value for a hormone could not be included in the analyses.

### Analyses

Linear mixed models were used to test for possible effects of hormonal status on disgust sensitivity. Analyses were conducted using R version 3.3.2 (R Core Team, 2016), with lme4 version 1.1-13 (Bates et al., 2014) and lmerTest version 2.0-33 (Kuznetsova et al., 2013). The dependent variable was Three Domain Disgust subscale score (separate models were run for each of the three disgust subscales). Predictors were the scaled and centered hormone levels. Random slopes were specified maximally following Barr et al. (2013) and Barr (2013). That is, random slopes were included for all within-woman predictors and, for analyses including interactions between different within-woman predictors (see Barr et al., 2013), the random slope for the interaction was included instead of the random slopes for each of the individual predictors (see Barr, 2013). Simulations have shown that models that do not include these random slopes have unacceptably high false positive rates (Barr et al., 2013; Barr, 2013). Full model specifications and full results for each analysis are given in our Supplemental Information. Data files and analysis scripts are publicly available at https://osf.io/93n2d/.

## Results

Scores for each Three Domain Disgust subscale were analyzed separately. For each subscale score we ran three models. Our first model (Model 1) included estradiol, progesterone, and their interaction as predictors. Our second model (Model 2) included estradiol, progesterone, and estradiol-to-progesterone ratio as predictors. Our third model (Model 3) included testosterone and cortisol as predictors, but did not consider possible effects of estradiol or progesterone. This analysis strategy is identical to that used in Jones et al. (2017a) and Jones et al. (2017b) to investigate the hormonal correlates of women’s face preferences and sexual desire, respectively. We adopted this analysis strategy because Model 1 closely follows the model used by Puts et al. (2013) to assess within-woman, fertility-linked effects. We include Model 2 because some research has used estradiol-to-progesterone ratio, rather than the interaction between estradiol and progesterone, to test for the combined effects of estradiol and progesterone (e.g., Eisenbruch et al., 2015). Model 3 is included because, although not typically considered in those models, testosterone and cortisol have been found to predict within-woman changes in behavior in other work (e.g., Ditzen et al., 2017; Welling et al., 2007). Thus, our models were chosen a priori to represent the variety of methods used in the literature on effects of hormone levels on women’s behavior.

### Pathogen disgust

Model 1 revealed no significant effects of progesterone (estimate=0.32, t=0.74, p=.46), estradiol (estimate=0.30, t=0.61, p=.55), or the interaction between estradiol and progesterone (estimate=0.22, t=0.09, p=.93). Model 2 also revealed no significant effects of progesterone (estimate=0.02, t=0.05, p=.96), estradiol (estimate=0.45, t=0.89, p=.38), or estradiol-to-progesterone ratio (estimate=-0.35, t=-1.04, p=.31). Model 3 showed no significant effects of either testosterone (estimate=0.35, t=0.71, p=.48) or cortisol (estimate=0.02, t=0.06, p=.95).

### Sexual disgust

Model 1 revealed no significant effects of progesterone (estimate=0.26, t=0.70, p=.48), estradiol (estimate=-0.16, t=-0.36, p=.72), or the interaction between estradiol and progesterone (estimate=1.01, t=0.47, p=.64). Model 2 also revealed no significant effects of progesterone (estimate=0.39, t=0.92, p=.36), estradiol (estimate=-0.19, t=-0.43, p=.67), or estradiol-to-progesterone ratio (estimate=0.11, t=0.49, p=.63). Model 3 showed no significant effects of either testosterone (estimate=0.07, t=0.15, p=.88) or cortisol (estimate=0.24, t=0.74, p=.46).

### Moral disgust

Model 1 revealed no significant effects of progesterone (estimate=0.34, t=0.71, p=0.48), estradiol (estimate=0.32, t=0.57, p=.57), or the interaction between estradiol and progesterone (estimate=2.39, t=0.86, p=.39). Model 2 also revealed no significant effects of progesterone (estimate=0.47, t=0.86, p=.39), estradiol (estimate=0.30, t=0.53, p=.60), or estradiol-to-progesterone ratio (estimate=0.07, t=0.24, p=.81). Model 3 showed no significant effects of either testosterone (estimate=0.98, t=1.73, p=.084) or cortisol (estimate=-0.92, t=-1.88, p=.064).

### Additional analyses

We conducted some additional exploratory analyses at the request of an anonymous reviewer. First, we repeated each of the analyses described above controlling for test session order. No significant hormonal effects were evident in these analyses. Second, we tested for a zero-order effect of progesterone on pathogen disgust (i.e., ran Model 1 with progesterone as the only predictor). This test showed no significant effect of progesterone. Third, we tested for a between-women progesterone-pathogen disgust correlation using data from each participant’s first test session only. This analysis also showed no significant association between progesterone and pathogen disgust. These analyses are reported in full in our Supplemental Information.

## Discussion

The current study presents the strongest test to date of the Compensatory Prophylaxis Hypothesis by examining correlations between changes in salivary progesterone and pathogen disgust. We found no evidence that pathogen disgust tracked changes in women’s salivary progesterone. By contrast with previous research (Fleischman and Fessler, 2011; Żelaźniewicz et al., 2016), our results show no support for the hypothesis that raised progesterone levels are associated with increased disgust responses to pathogen cues (Fessler et al., 2005; Fleischman & Fessler, 2011). We also found no evidence that pathogen disgust tracked changes in estradiol, testosterone, or cortisol.

Fessler and Naverrete (2003) reported that sexual disgust increased during the high-fertility phase of the menstrual cycle. They hypothesized that this hormone-linked change in sexual disgust functioned to reduce the likelihood of sexual behaviors that could harm a woman’s reproductive fitness. By contrast with Fessler and Naverrete (2003), we found no evidence that sexual disgust tracked changes in women’s hormonal status, including changes that are highly correlated with fertility (e.g., changes in estradiol-to-progesterone ratio, Gangestad et al., 2016). Recent research has raised questions about the robustness of some hypothesized links between aspects of women’s hormonal status and mating psychology (see, e.g., Gangestad et al., 2016 for a discussion of some of these questions). Our null results for sexual disgust raise similar questions about the robustness of hypothesized links between women’s hormonal status and an aspect of mating psychology (sexual disgust) that had not yet been reassessed in the context of this discussion.

We believe that the current study provides the best test to date of the Compensatory Prophylaxis Hypothesis, for multiple reasons. First, we measured changes in both progesterone and disgust sensitivity within women over multiple observations. Second, our sample size was approximately four times larger than that used in earlier compensatory prophylaxis work (Fleischman & Fessler, 2011). Furthermore, although our work relied upon self-report, earlier work reporting support for the Compensatory Prophylaxis Hypothesis also used self-report (e.g., Fleischman & Fessler, 2011). That said, studies using psychophysiological measures of disgust sensitivity (see, e.g., De Smet et al., 2014) could yet reveal hormone-linked changes in disgust sensitivity that are not evident in analyses of self-report measures.

Whereas we administered the Three Domain Disgust Scale at multiple time point, some of the items administered by Fleischman and Fessler (2011) asked participants about pathogen avoidance specifically within the past 24 hours. Such item phrasing might be more sensitive to day-to-day fluctuations compared to the Three Domain Disgust Scale. That said, Fleischman and Fessler (2011) also asked participants to report disgust toward visual cues to pathogens, with a response format similar to that used in the current study. They reported the same support for the Compensatory Prophylaxis Hypothesis using this response format that did not mention behavior over the past 24 hours.

In conclusion, our results provide no support for the Compensatory Prophylaxis Hypothesis of pathogen disgust. We also found no evidence that sexual disgust tracks changes in women’s hormonal status. These results underline the importance of employing longitudinal designs, hormone measurements, and large samples to investigate hypothesized links between hormonal status and emotional responses.

## Open Practices

Data files and analysis scripts are publicly available via Open Science Framework at https://osf.io/93n2d/.

## Process Data

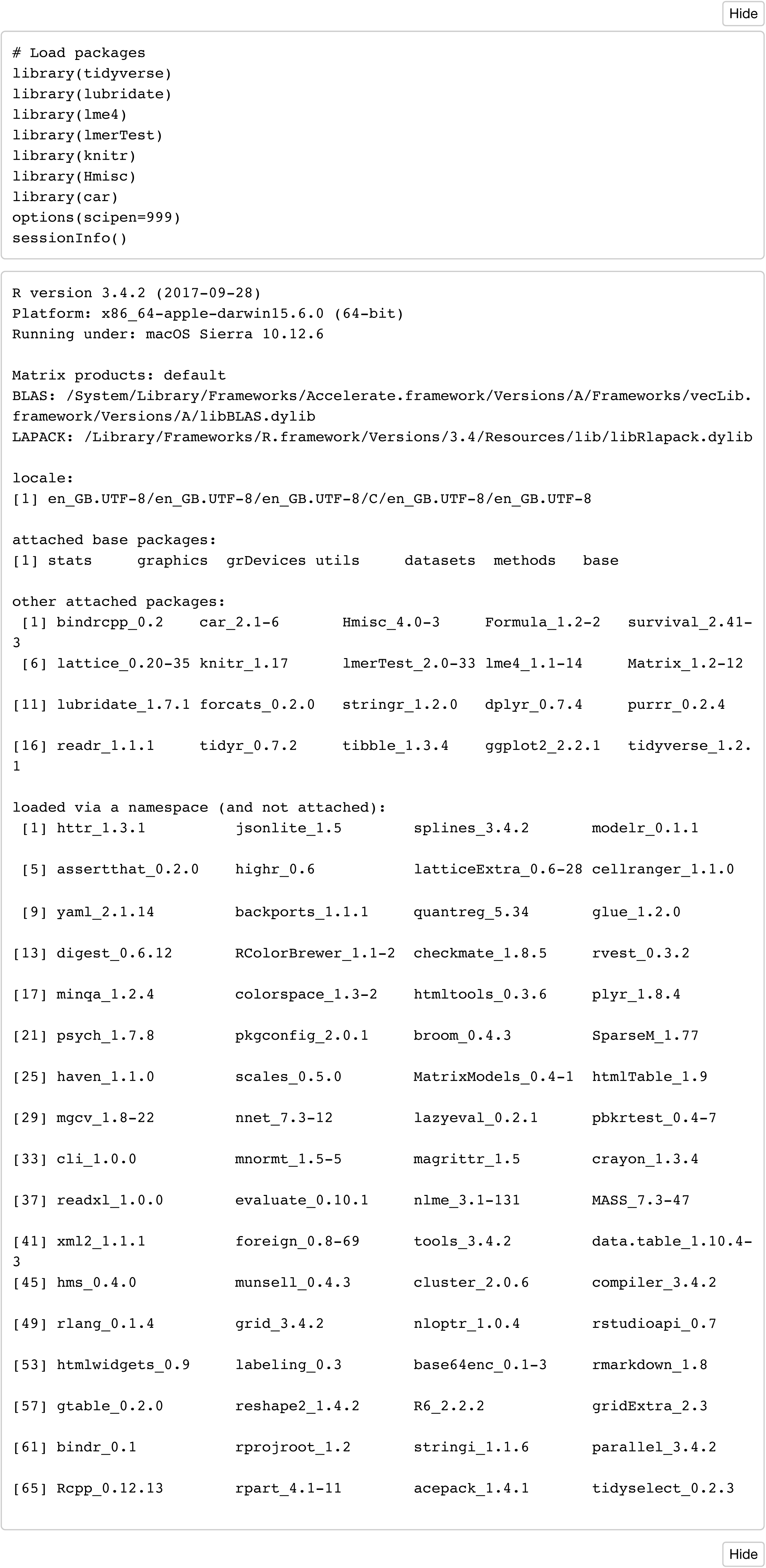

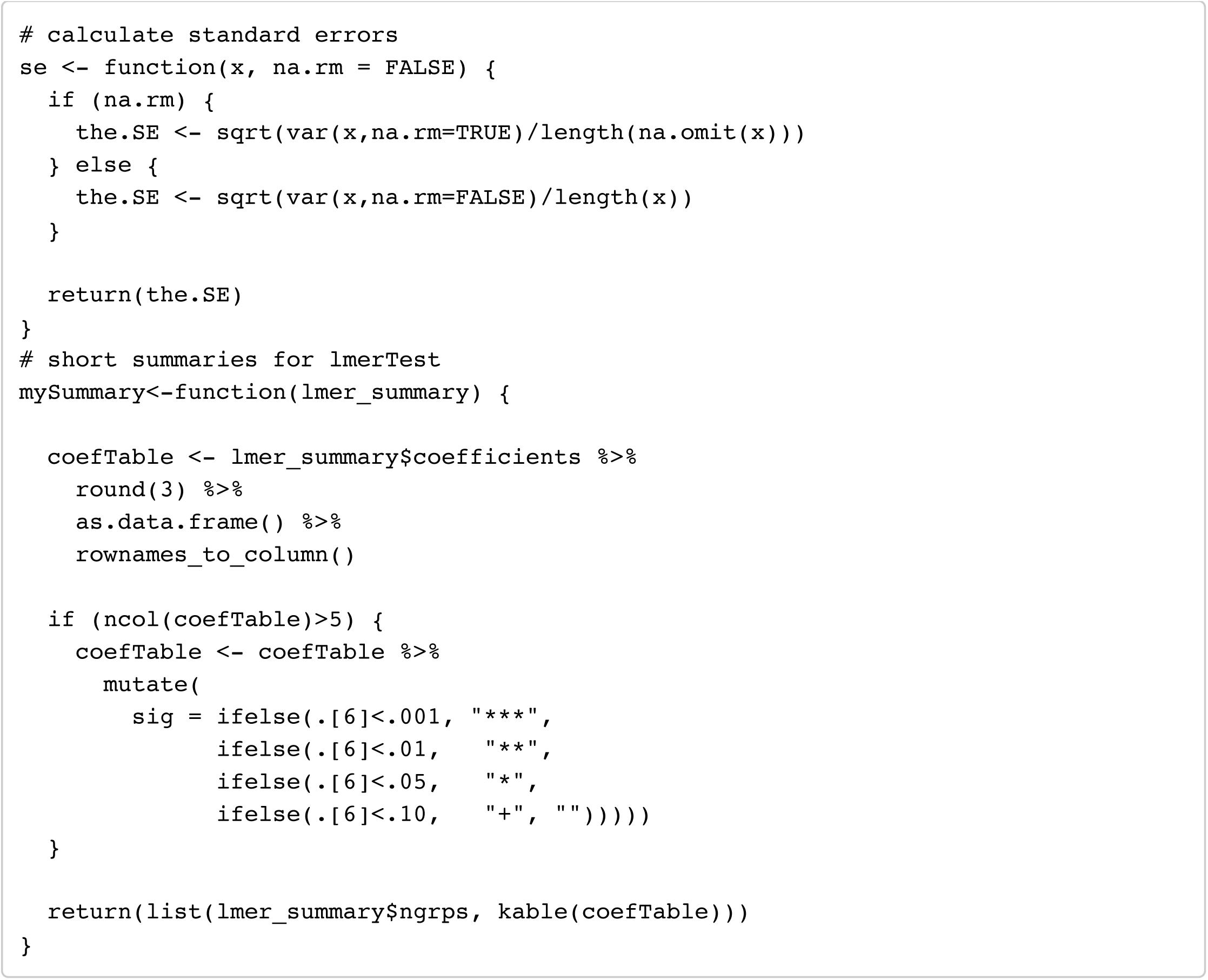

### Load full dataset

Data entered from all white, heterosexual women not using hormonal contraceptives. Each row is all data from a single session (i.e. oc_id:date)

- “oc_id” = ID of the subject
- “block” = testing block (1, 2 or 3)
- “age” = age (in years) of subject on day of testing
- “ethnicity” = ethnic group of subject (all white)
- “sexpref” = sexual preference of subject (all heterosexual)
- “date” = date of testing session
- “pill” = whether subject was using hormonal contraceptives (all 0)
- “block_N” = how many testing sessions completed in that block
- “prog” = salivary progesterone for that session
- “estr” = salivary estradiol for that session
- “test” = salivary testosterone for that session
- “cort” = salivary cortisol for that session
- “manip” = TDDS subscale: pathogen_disgust, sexual_disgust, moral_disgust
- “rating” = score on the TDDS subscale

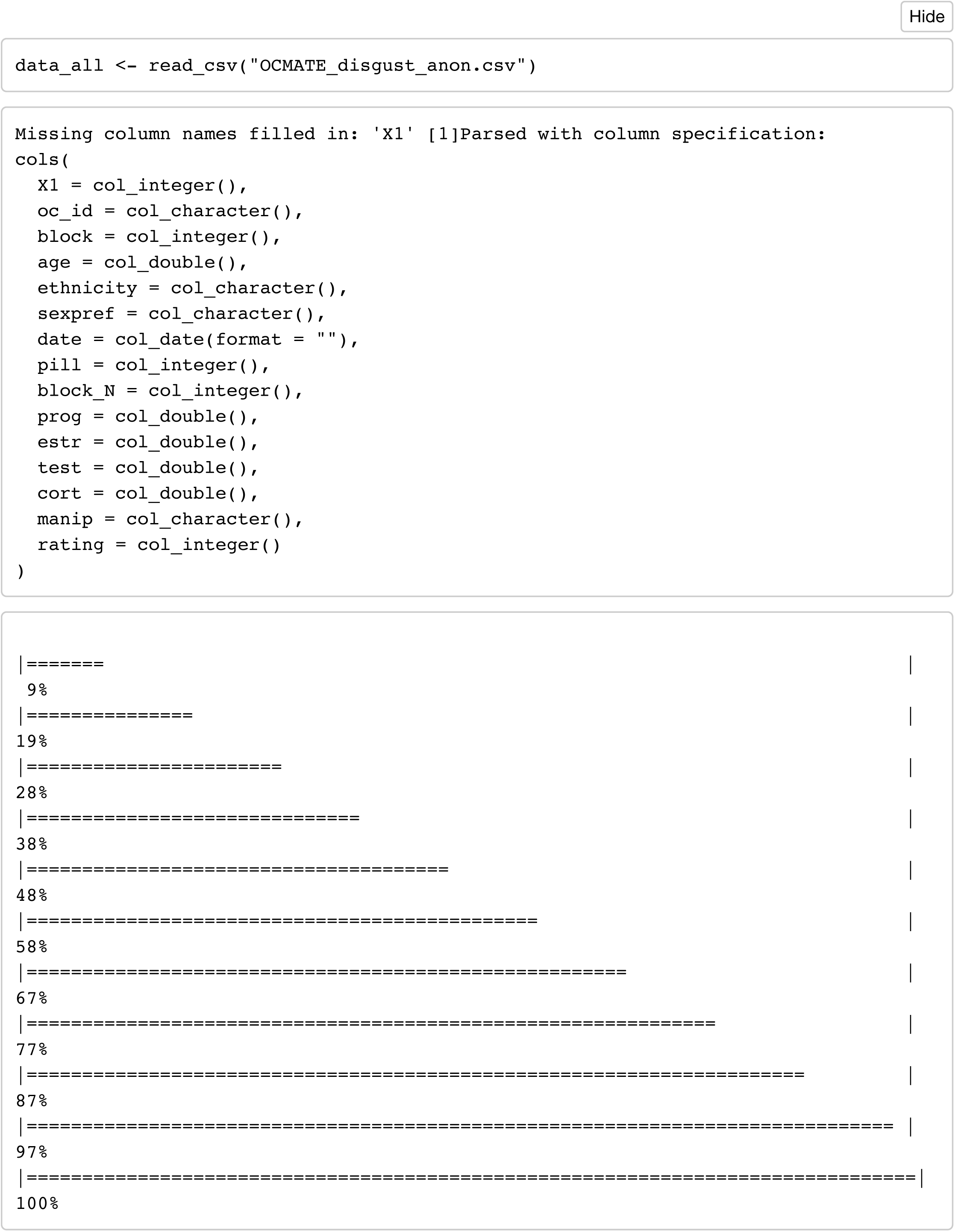

### Descriptive Stats

#### Age

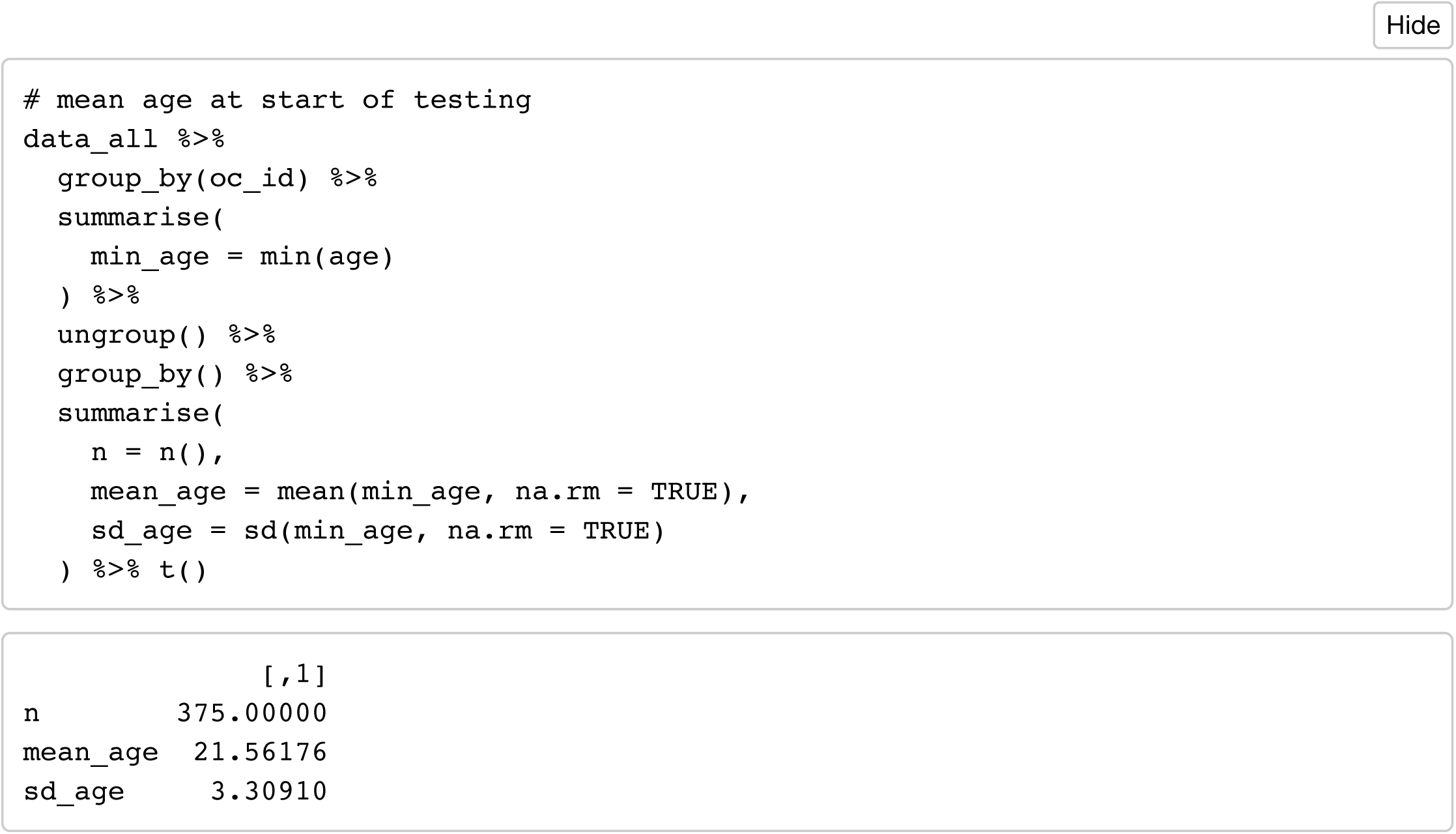

#### The number of sessions completed per woman

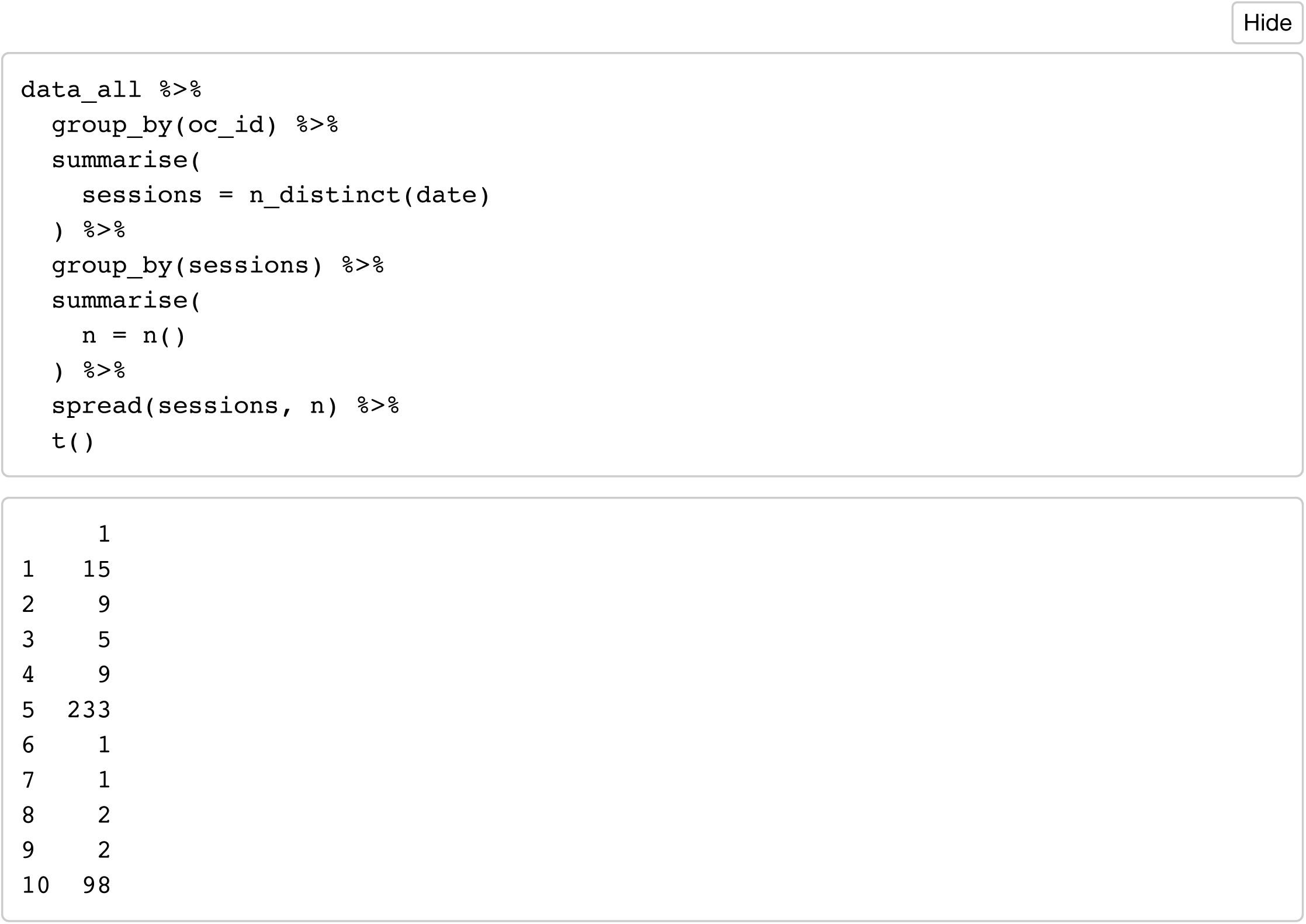

#### Block Intervals

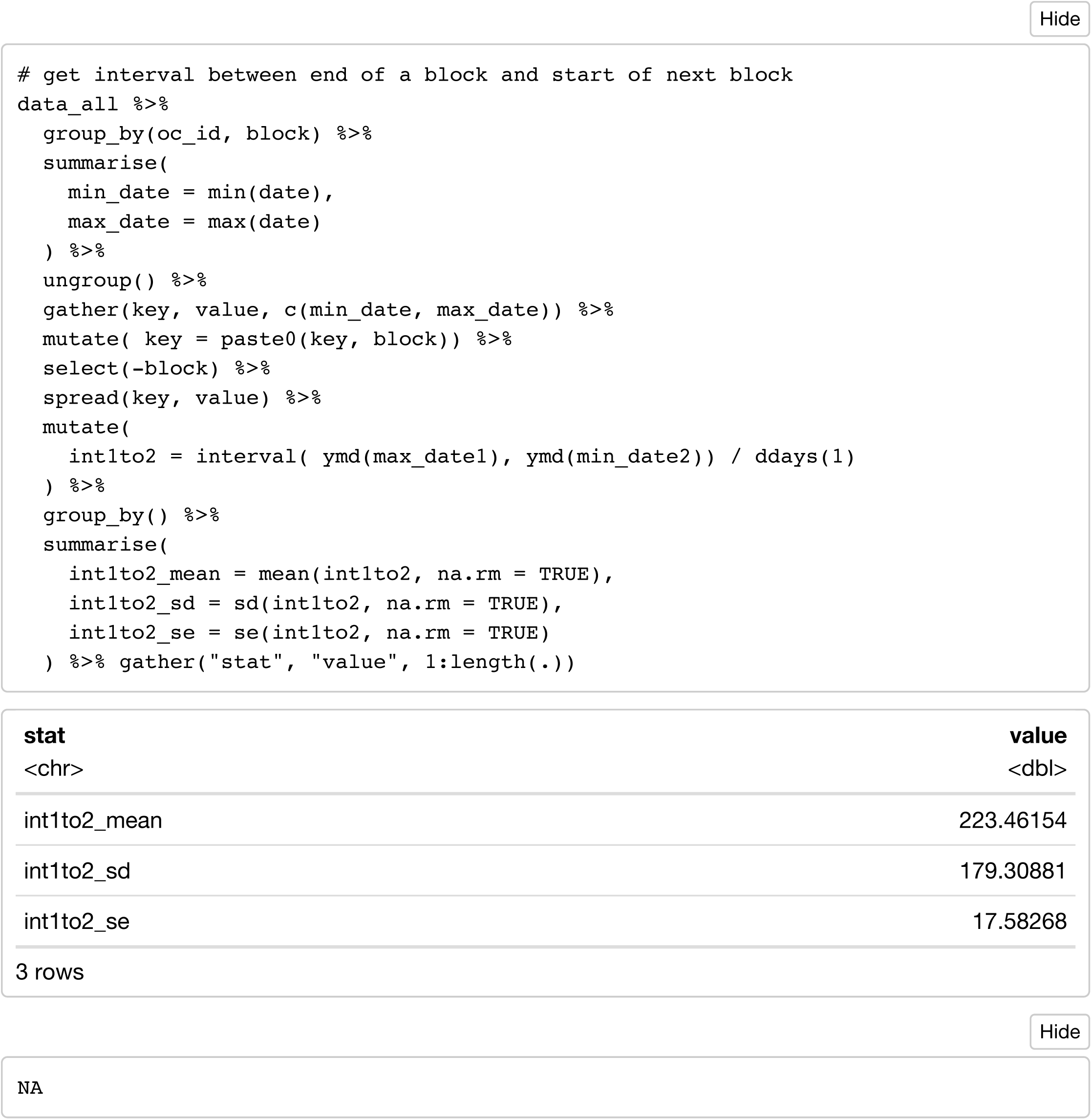

### Exclusions

#### Exclude observations with missing estr, prog and test

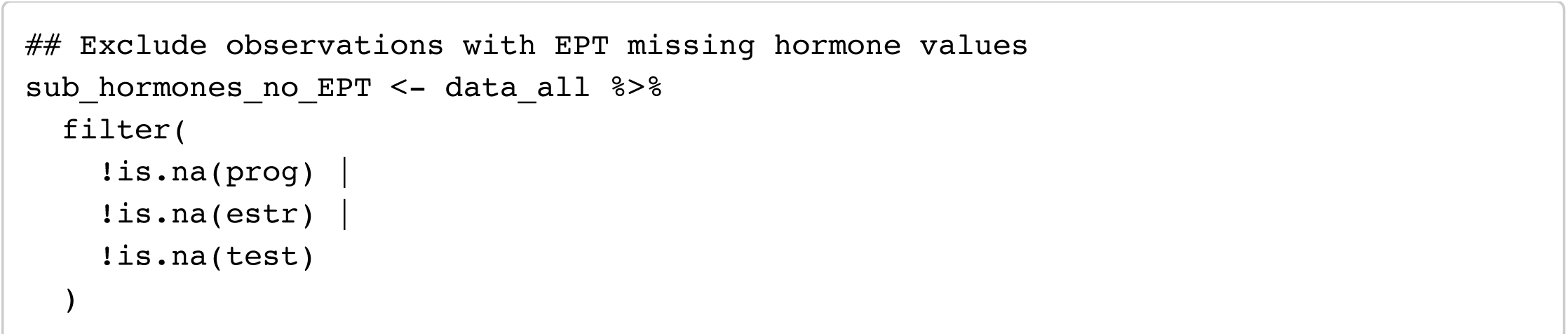

#### Exclude subjects with only a single session in a block

This is necessary because you can’t calculate subject-centered means with only one data point.

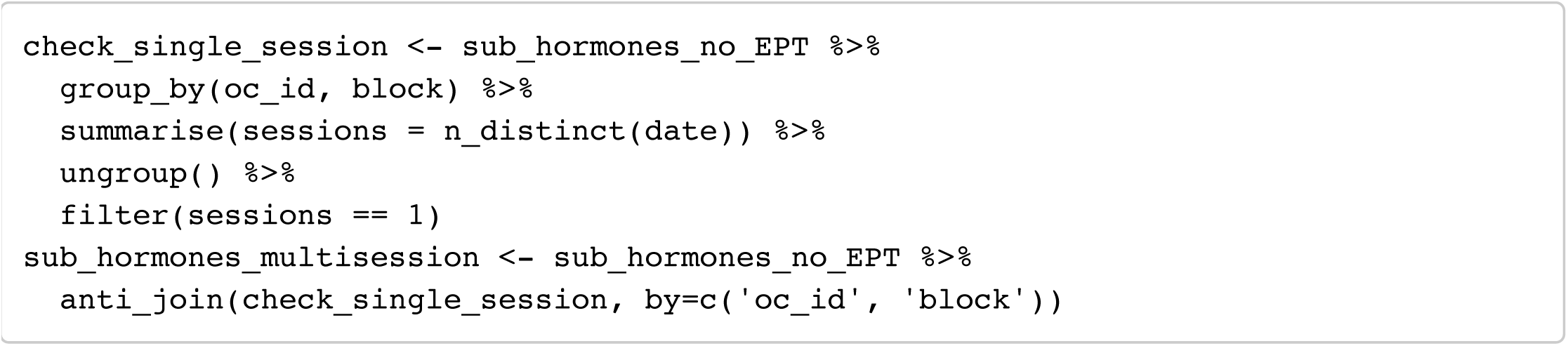

#### Remove outlier hormone values

Remove below bottom sensitivity thresholds for assays (progesterone < 5, estrogen < 0.1), and remove outlier values (+/− 3SD from the mean)

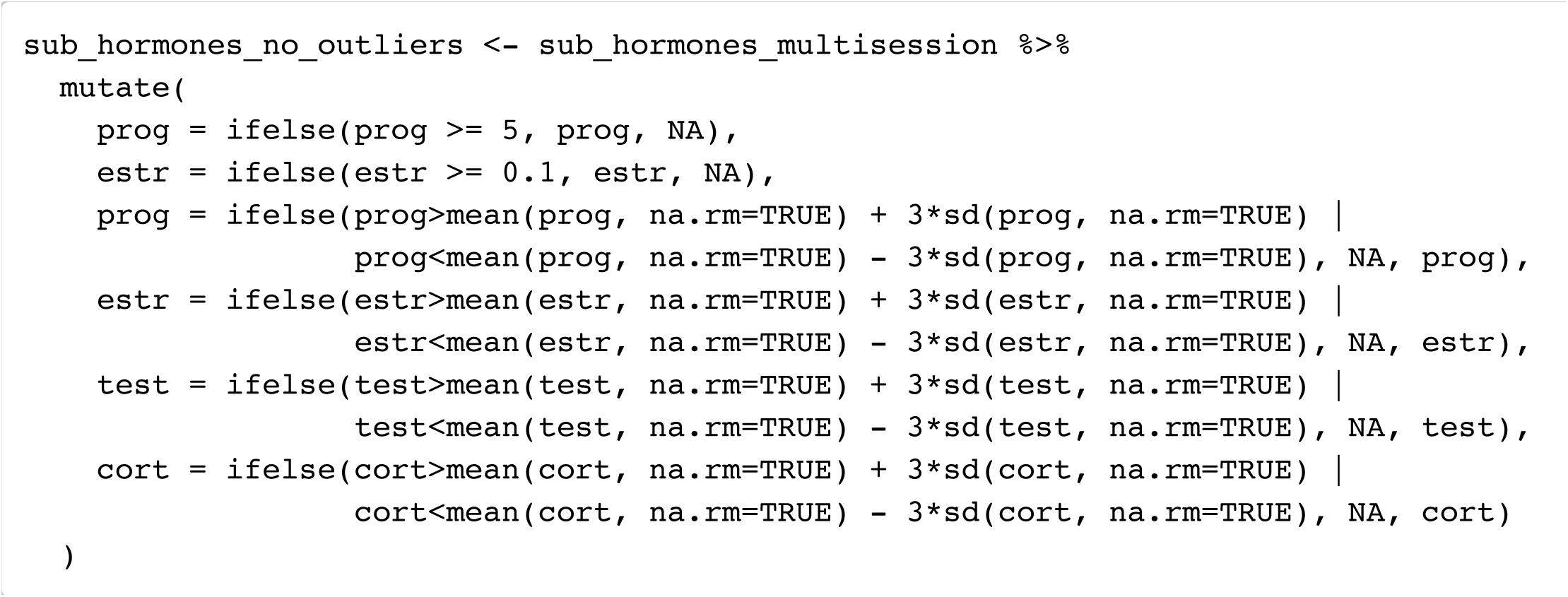

#### Numbers excluded and included

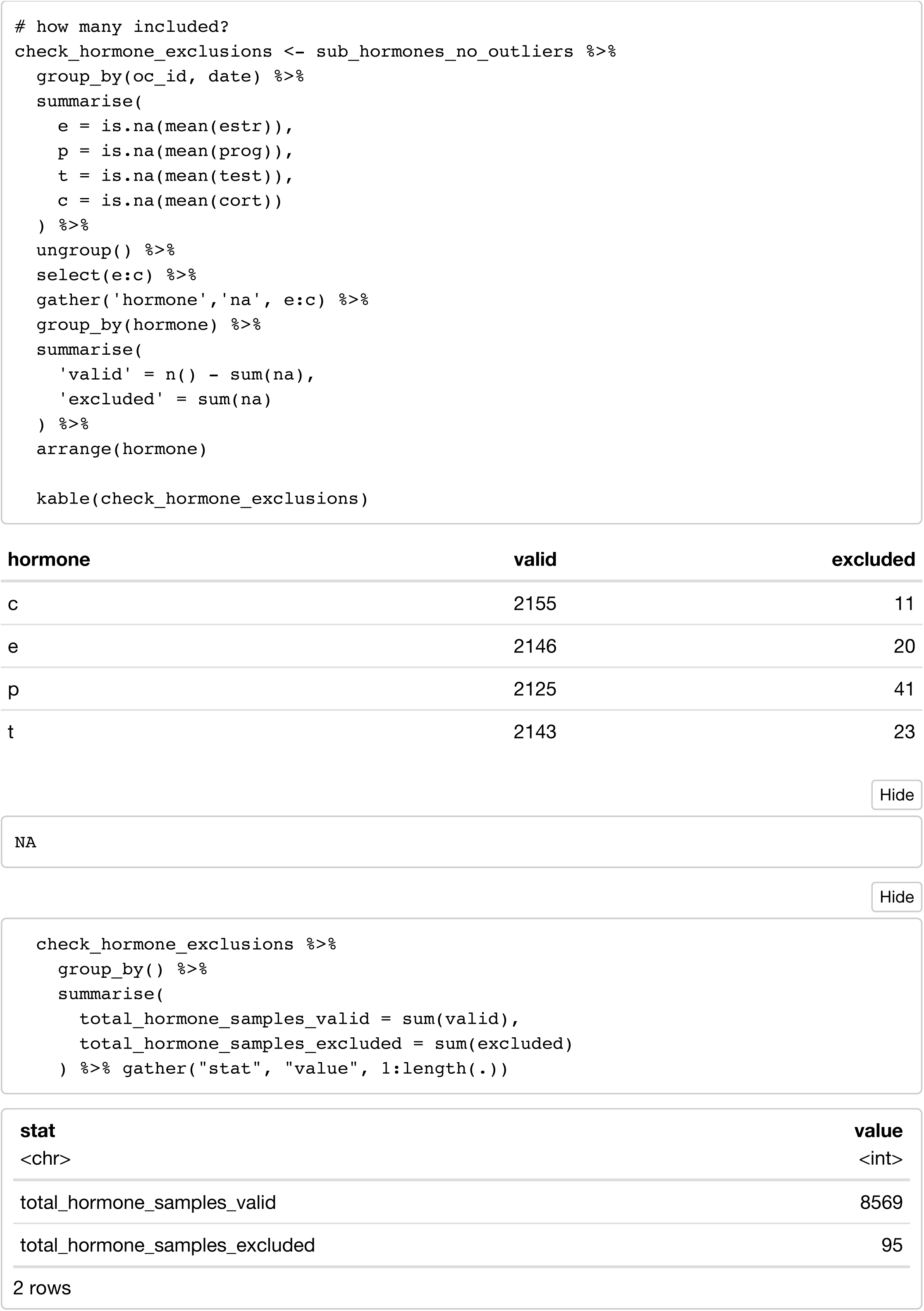

#### Subject-mean-centre hormones

Divide results by a constant to put all hormones on ~ −0.5 to +0.5 scale

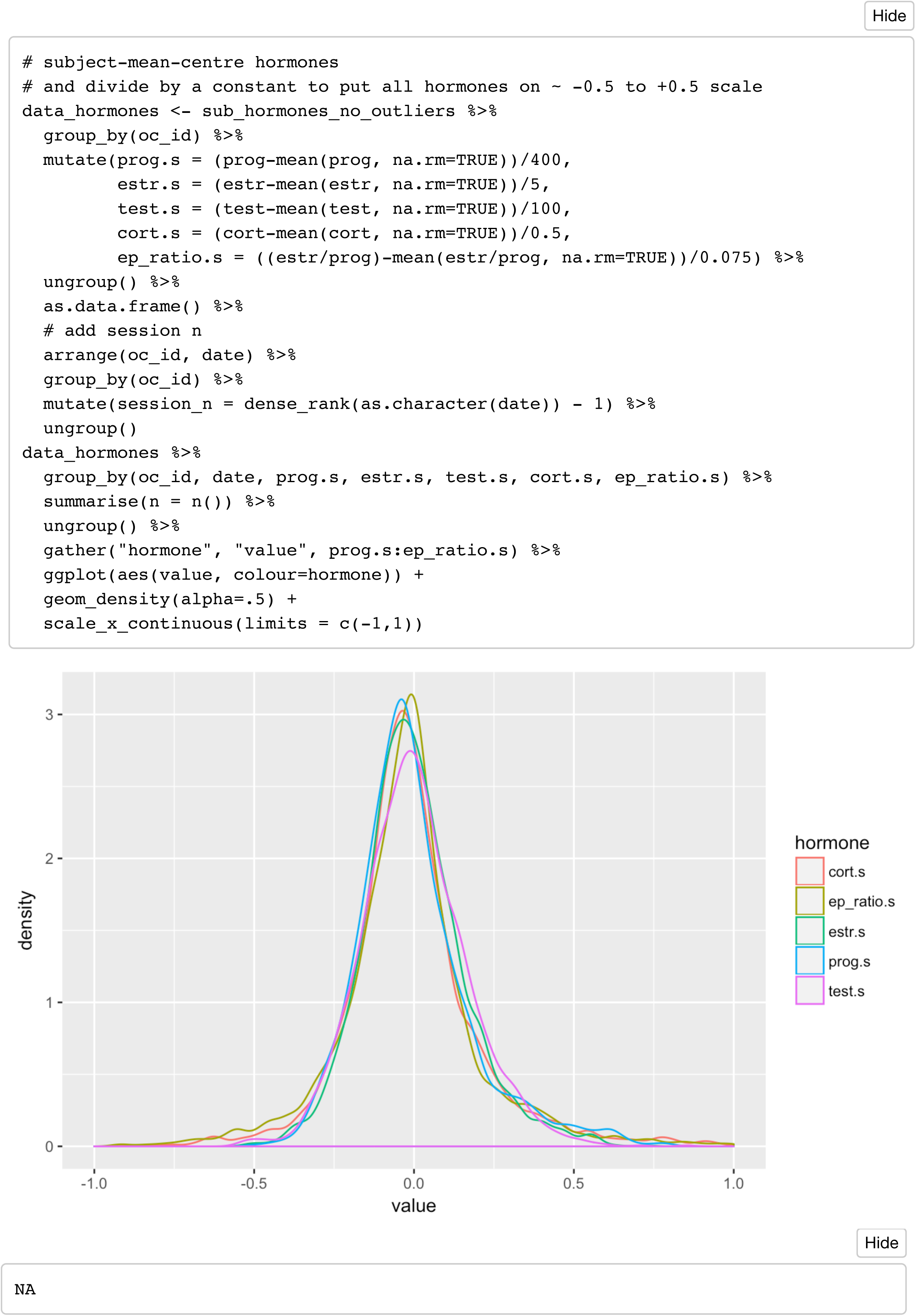

#### Mean Hormone Levels

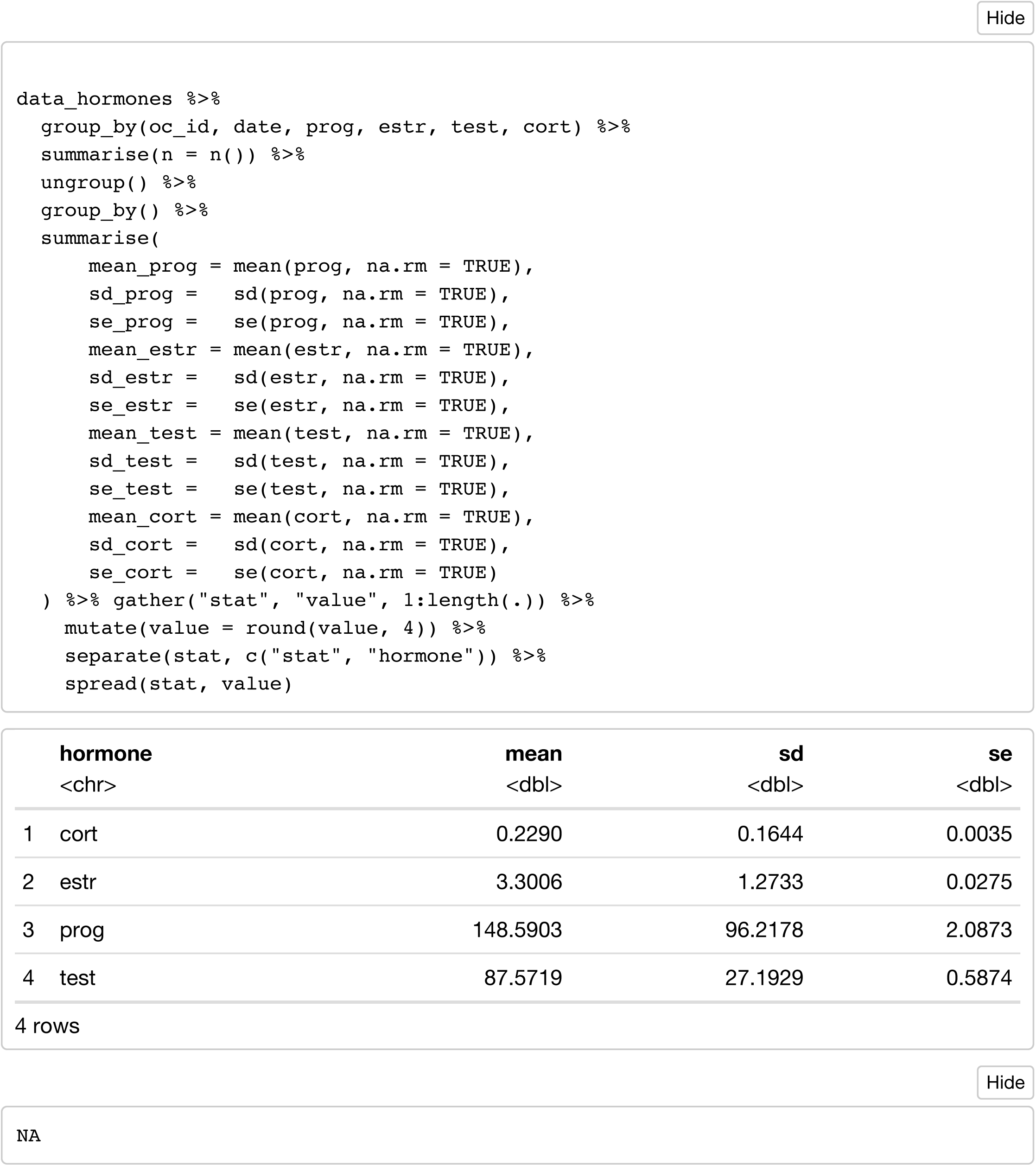

#### Mean hormone ranges

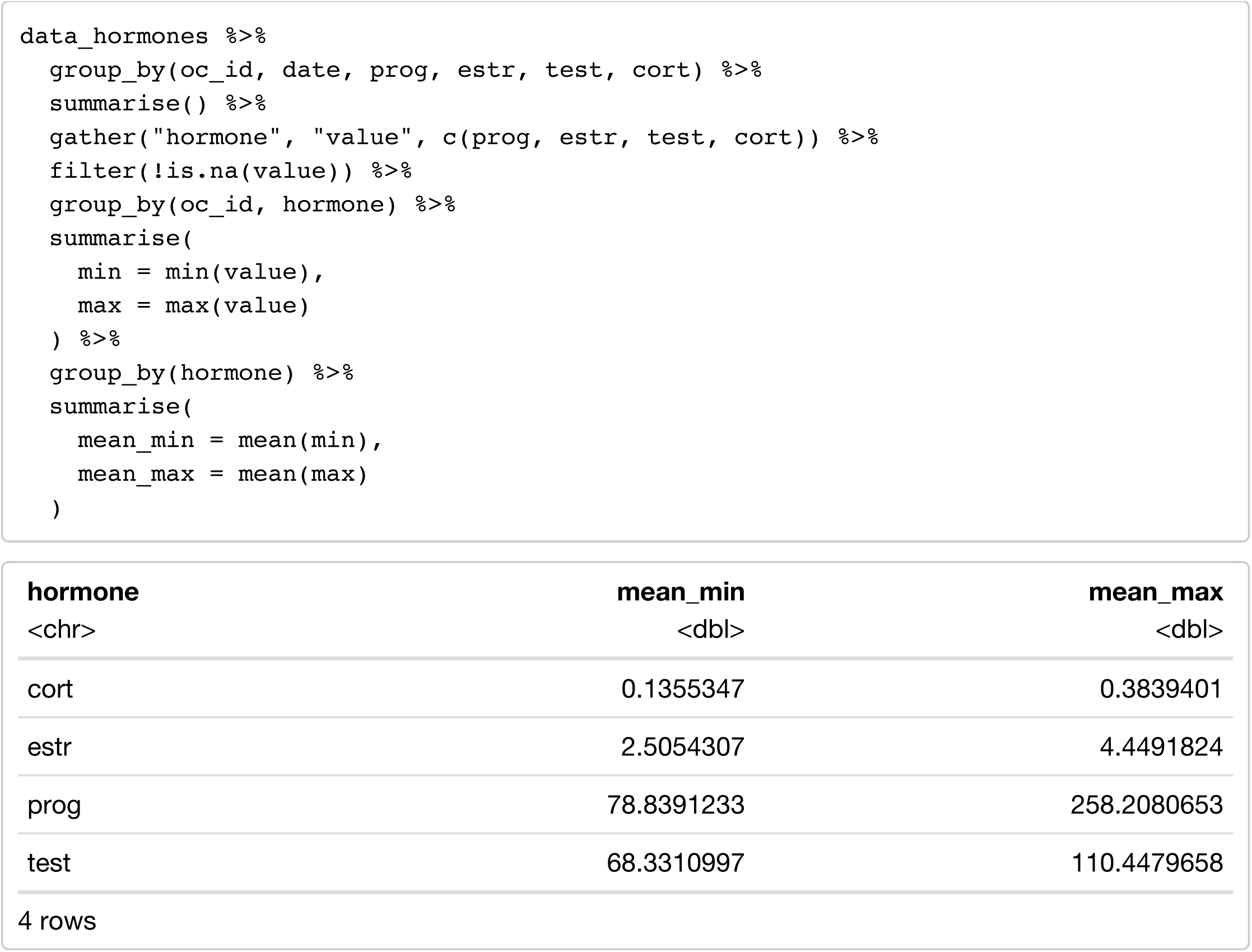

#### Descriptive Stats for moral, pathogen, and sexual disgust

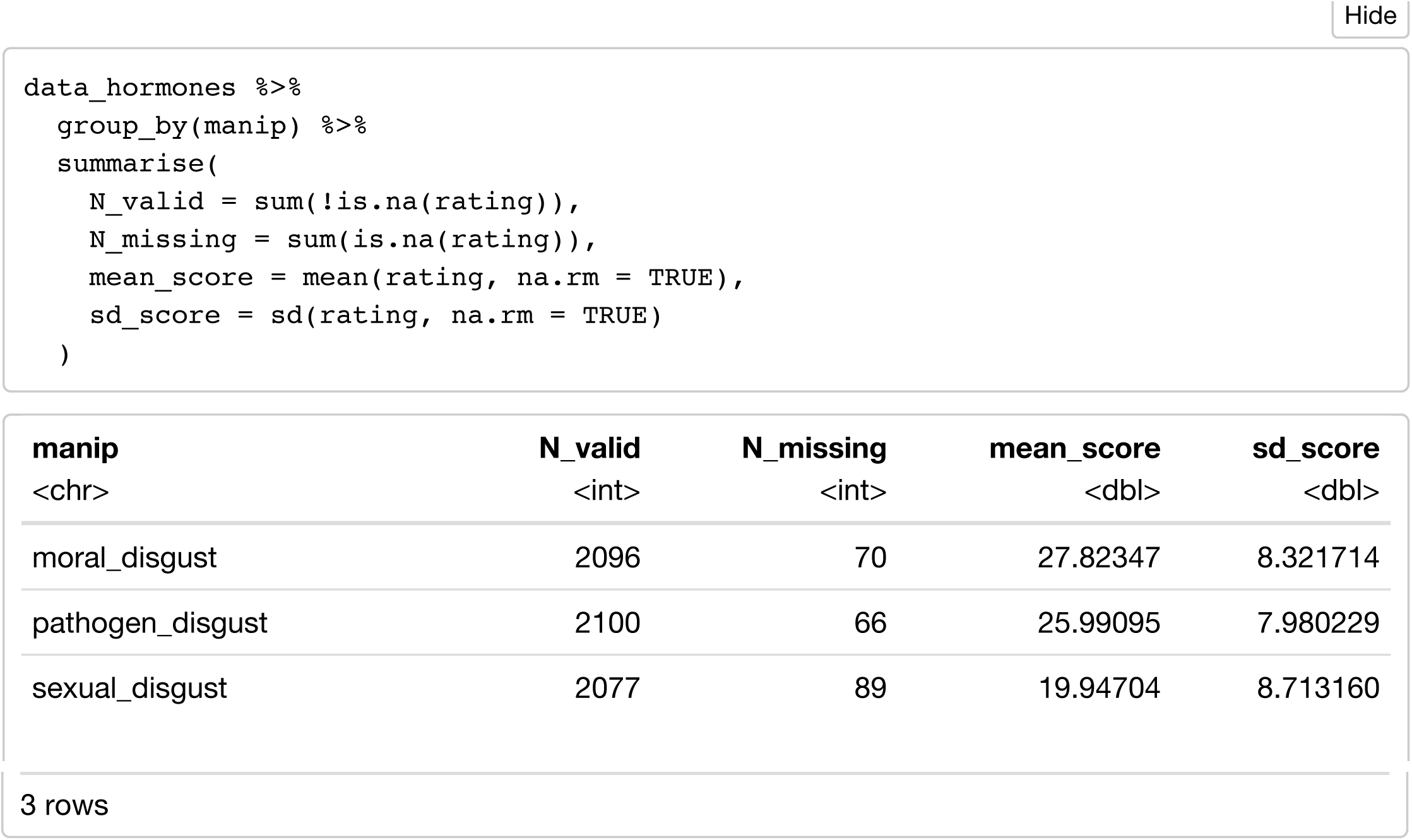

#### Relationship between P and Pathogen Disgust

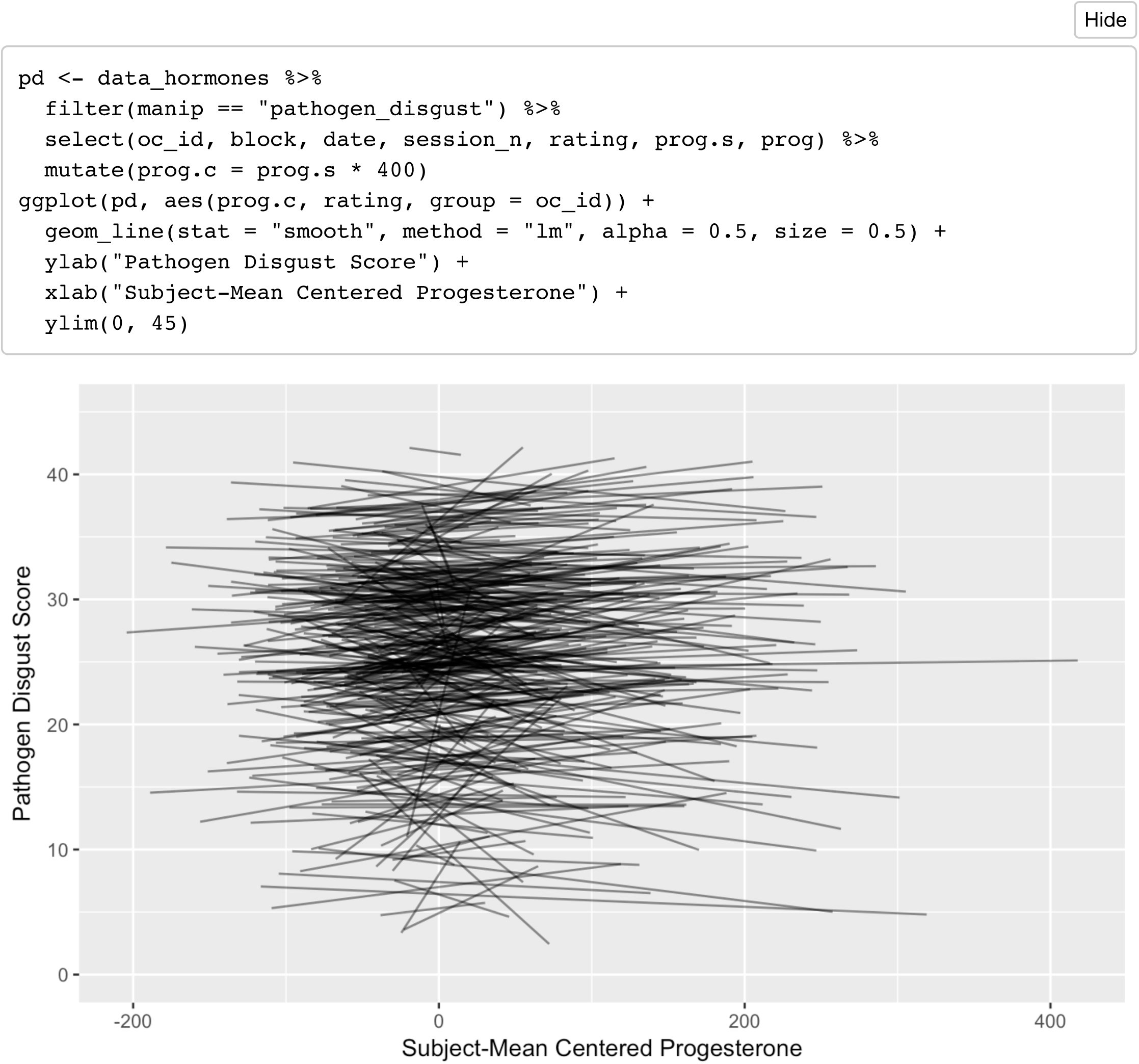

### Pathogen Disgust

#### Model 1: pathogen_disgust ~ E + P + E x P

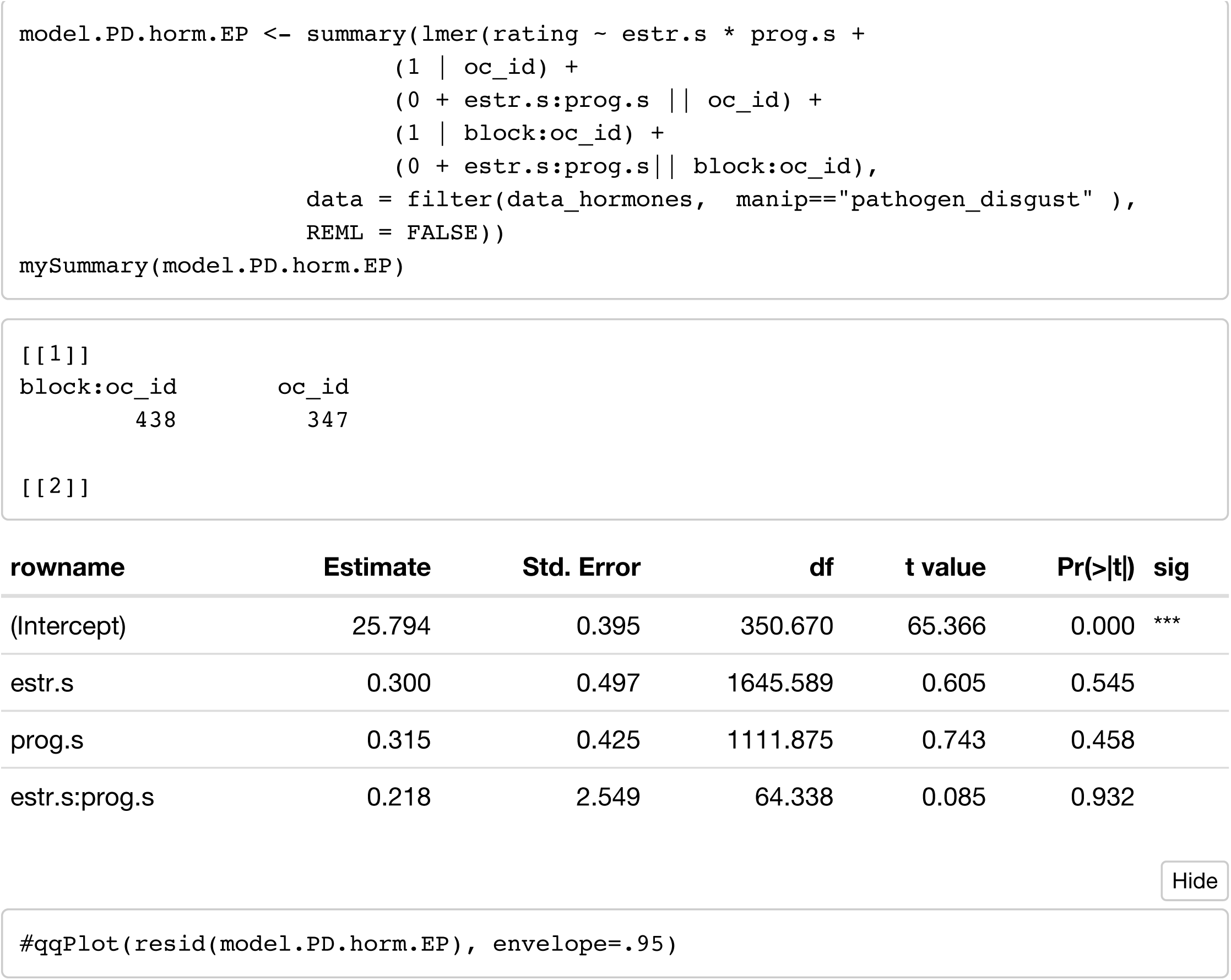

#### Model 2: pathogen_disgust ~ E + P + EP_ratio

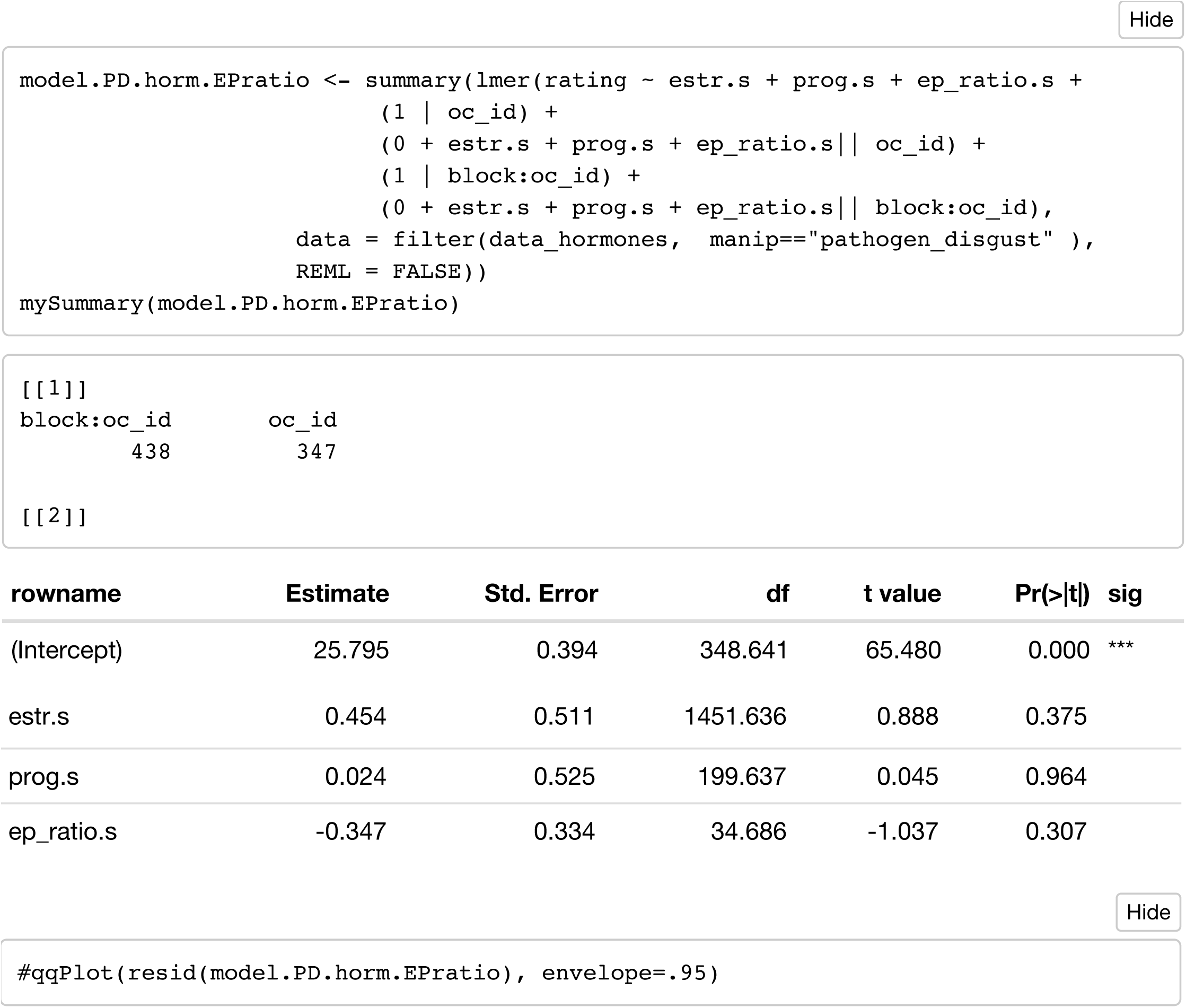

#### Model 3: pathogen_disgust ~ T + C

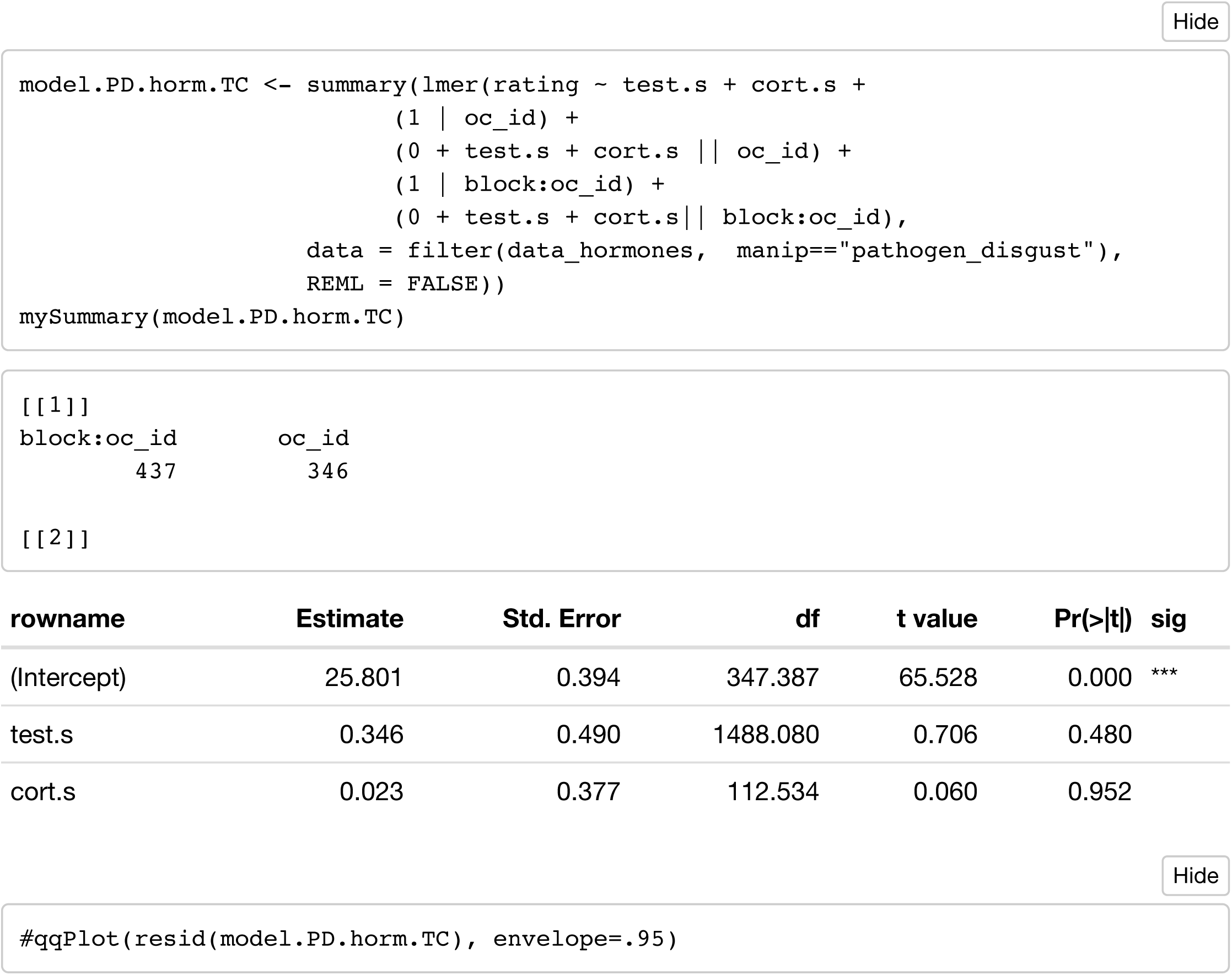

#### Model 4: pathogen_disgust ~ P

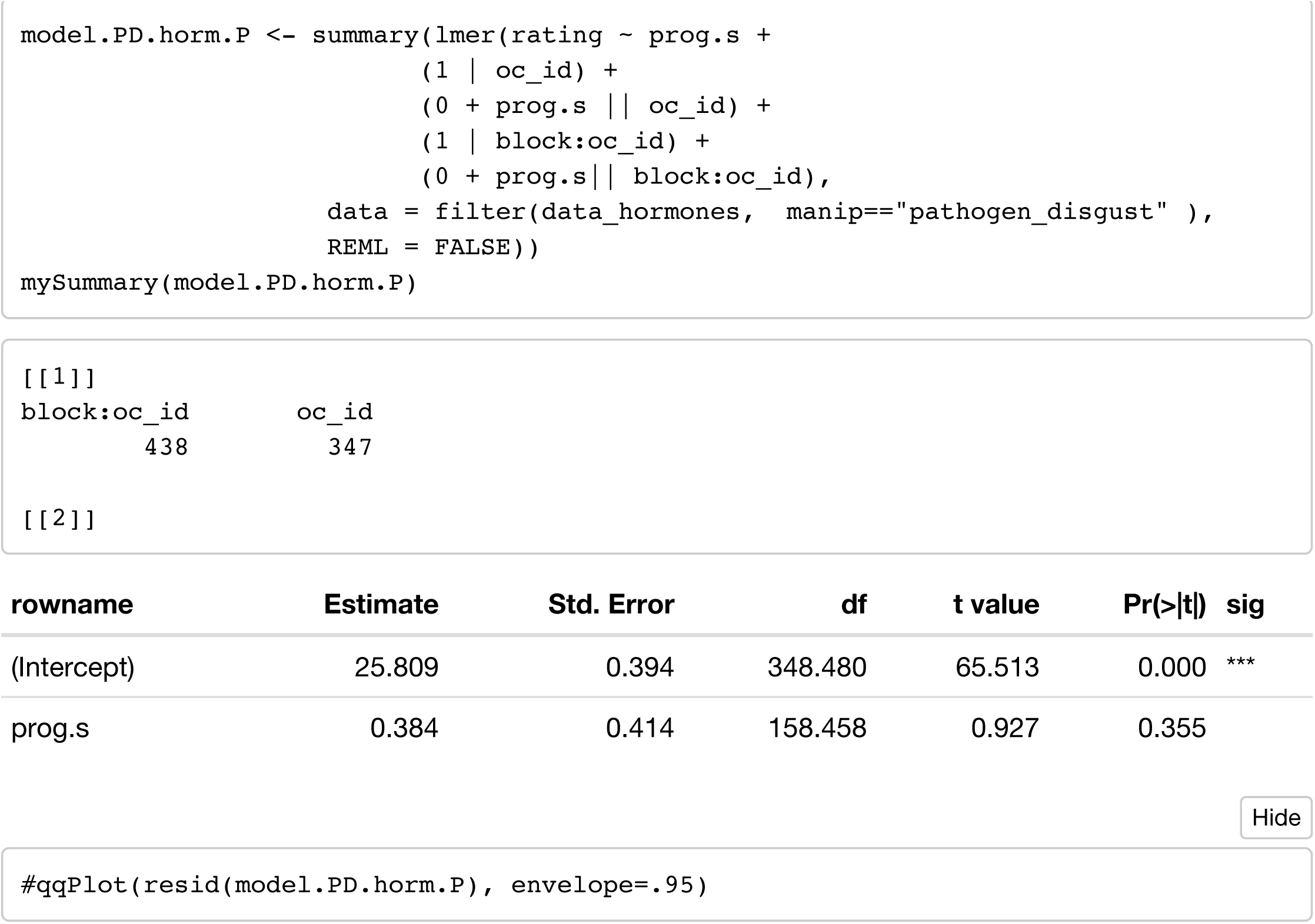

### Sexual Disgust

#### Model 1: sexual_disgust ~ E + P + E x P

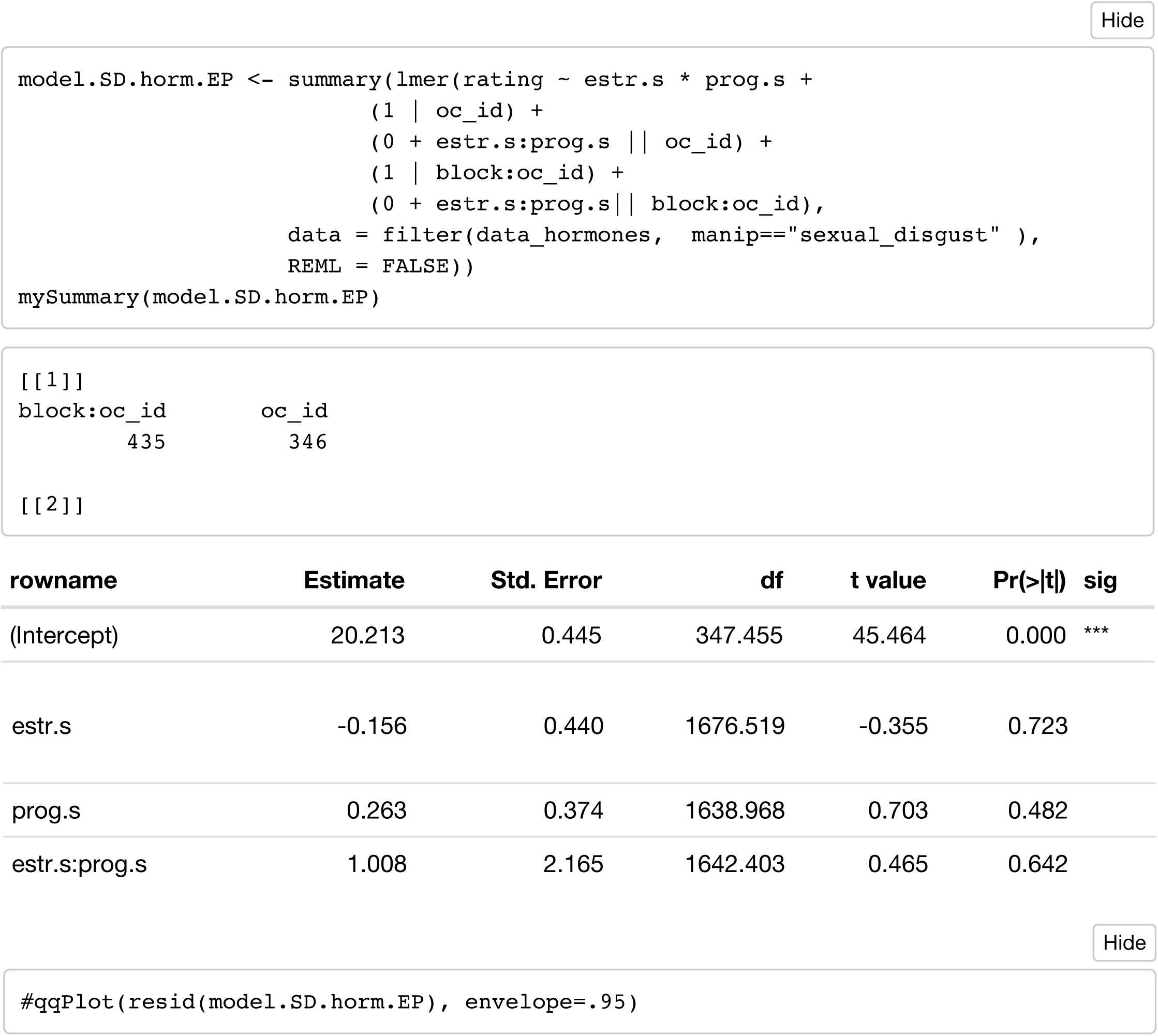

#### Model 2: sexual_disgust ~ E + P + EP_ratio

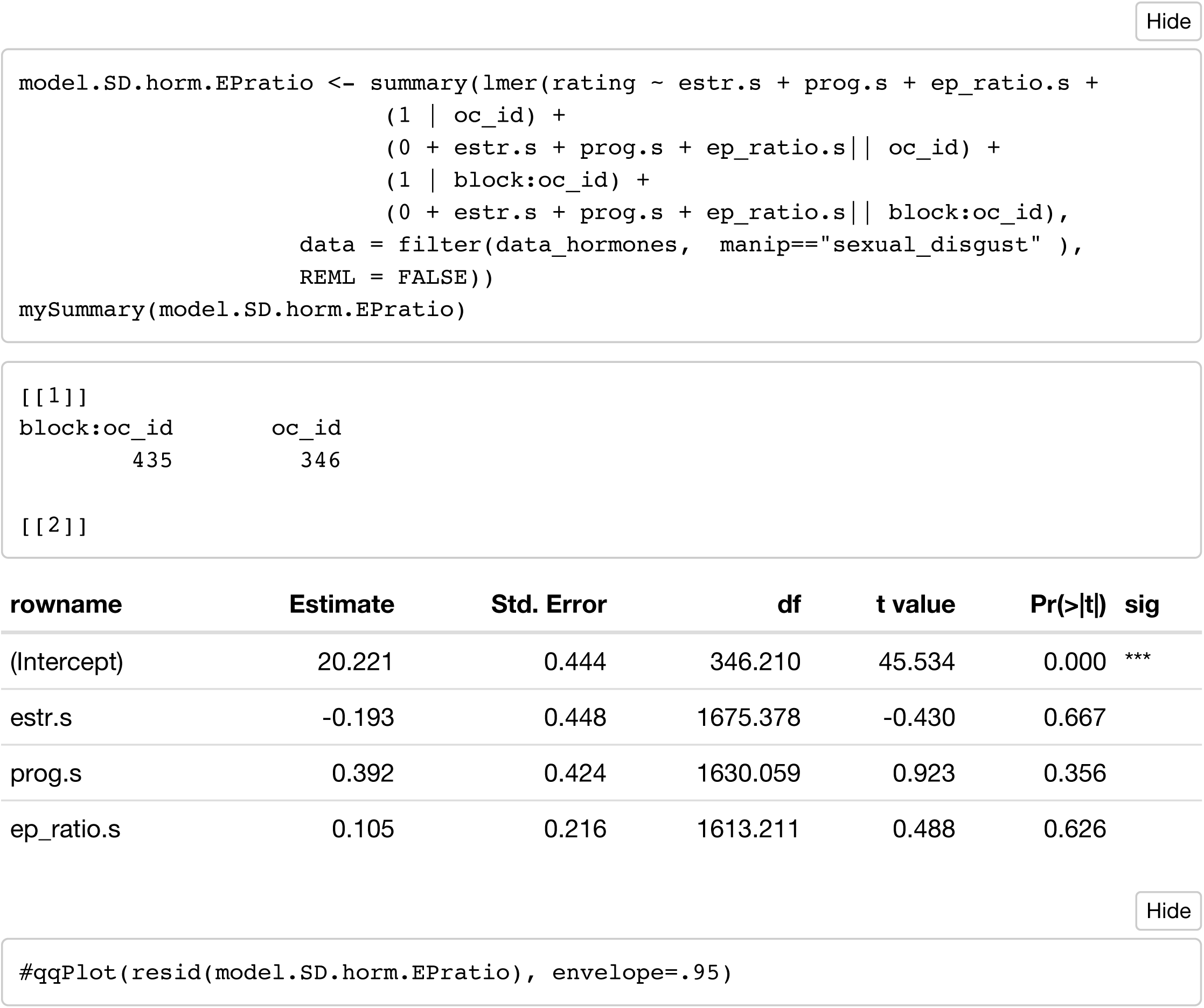

#### Model 3: sexual_disgust ~ T + C

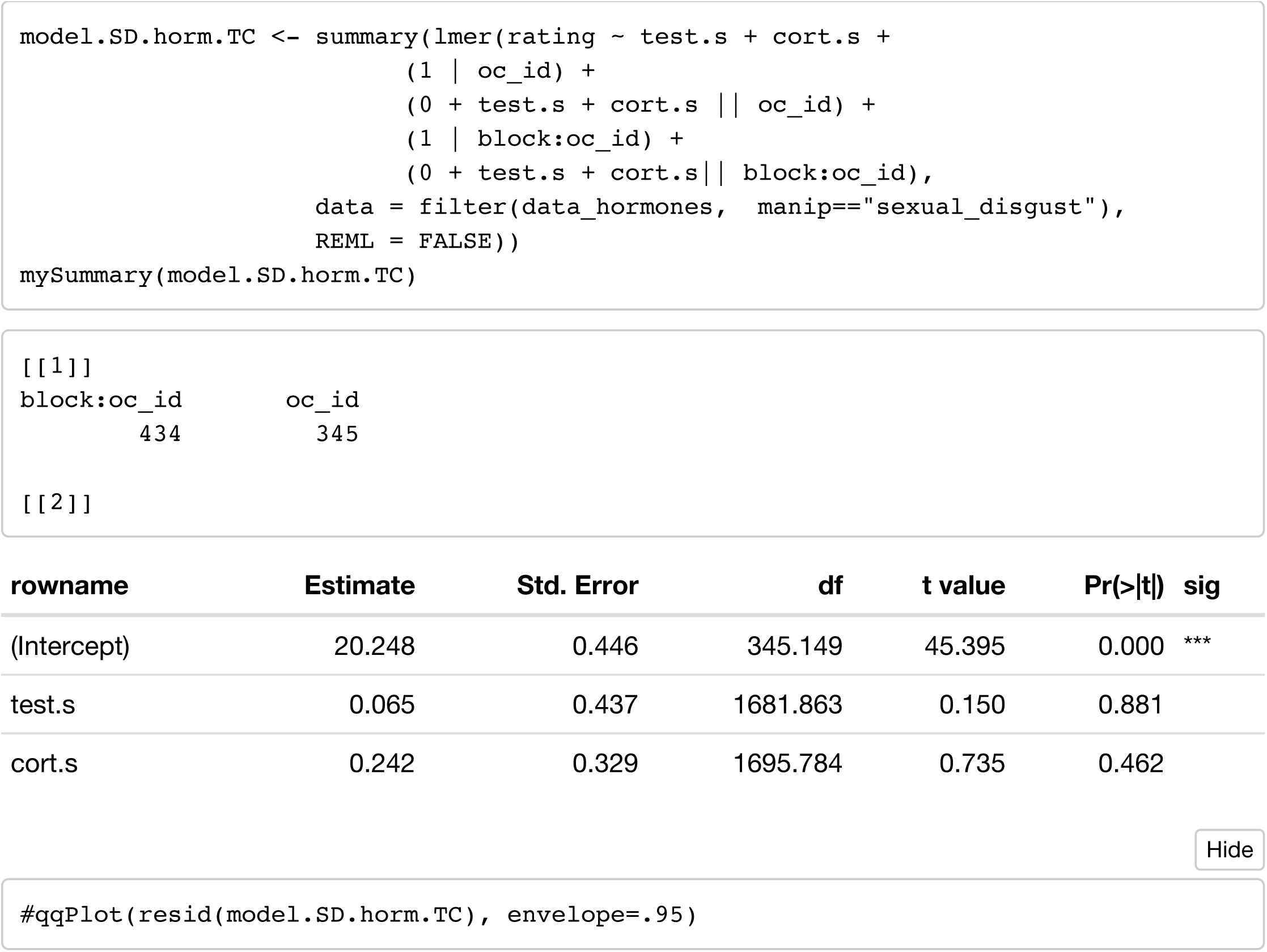

### Moral disgust

#### Model 1: moral_disgust ~ E + P + E x P

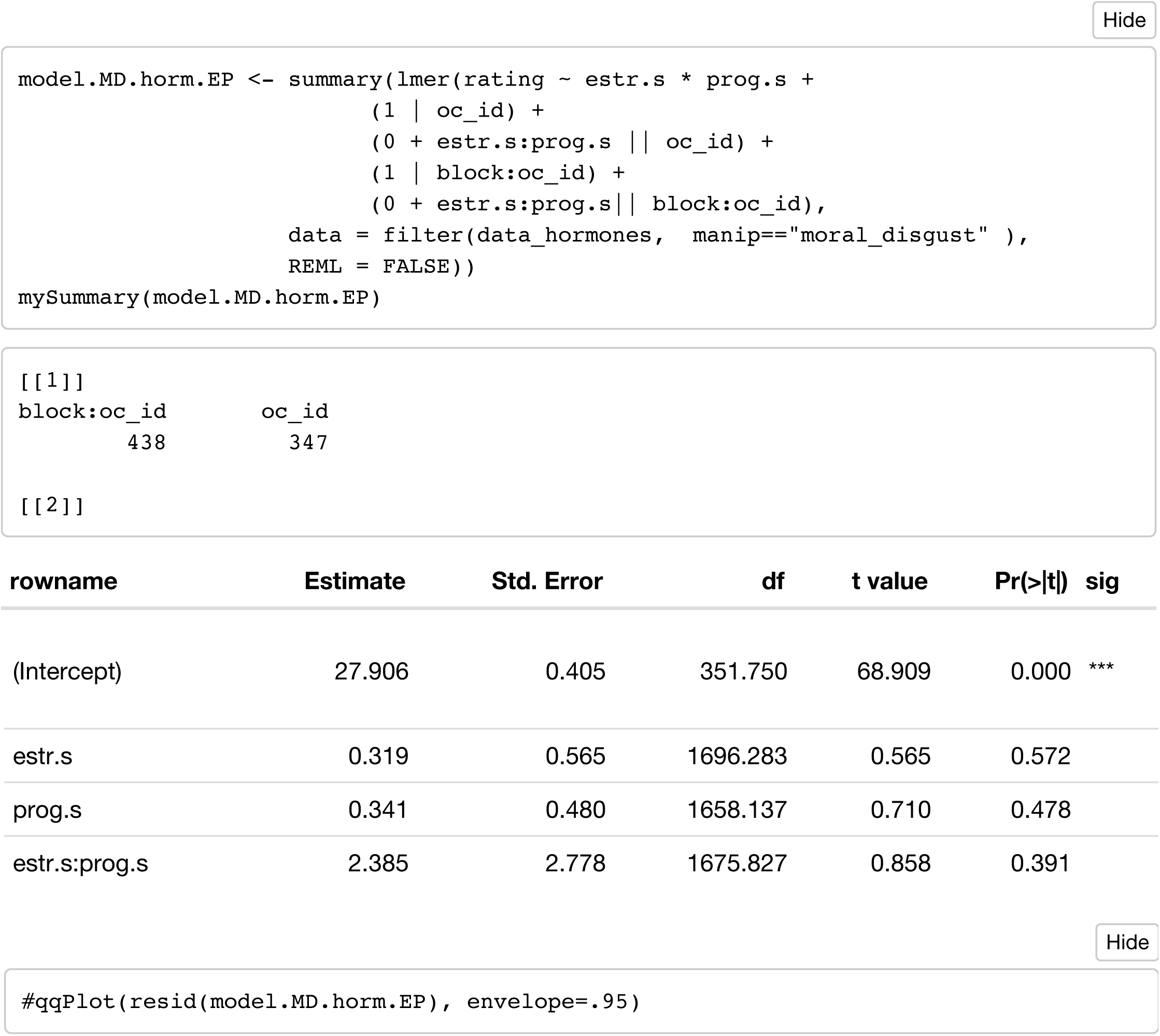

#### Model 2: moral_disgust ~ E + P + EP_ratio

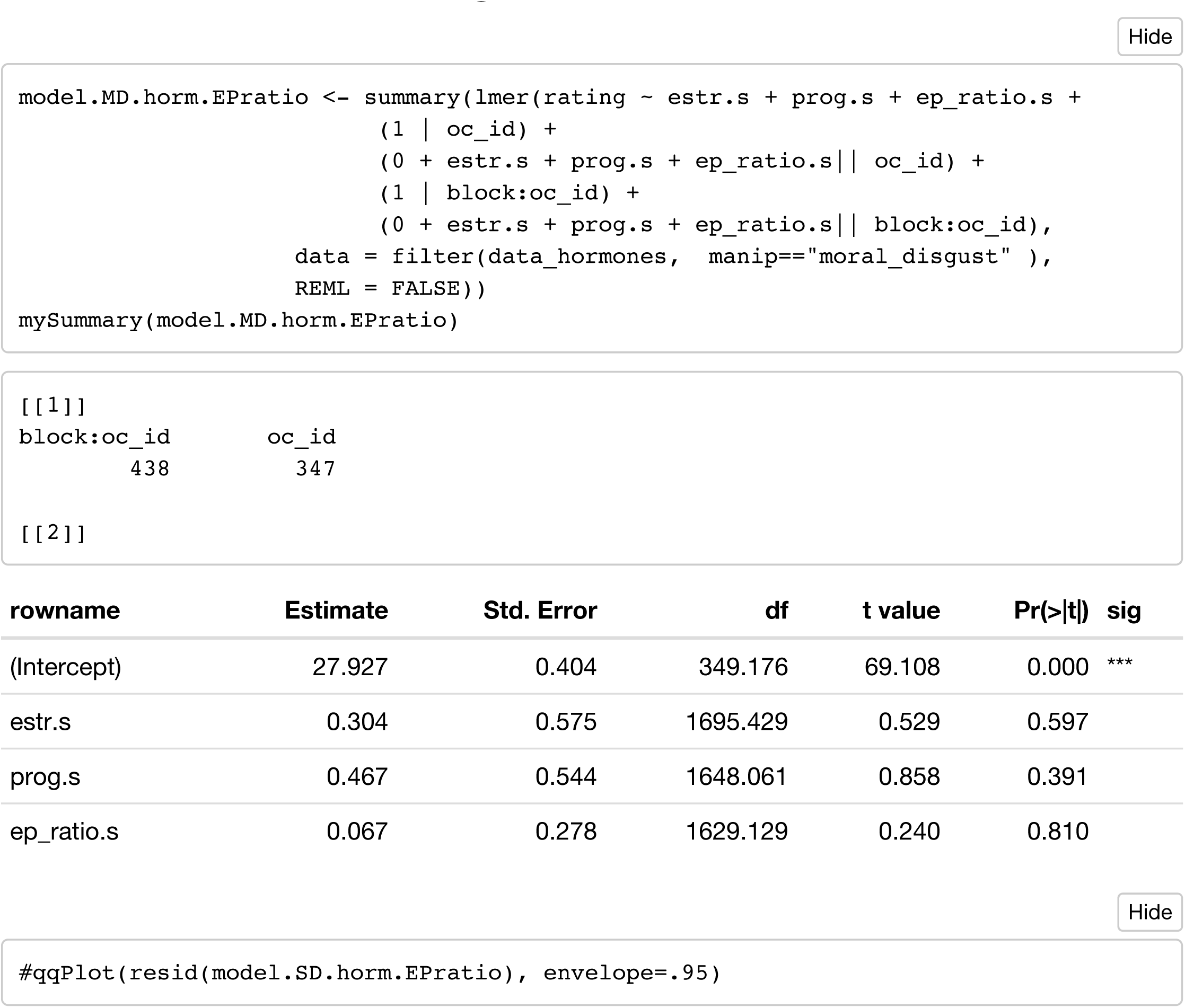

#### Model 3: moral_disgust ~ T + C

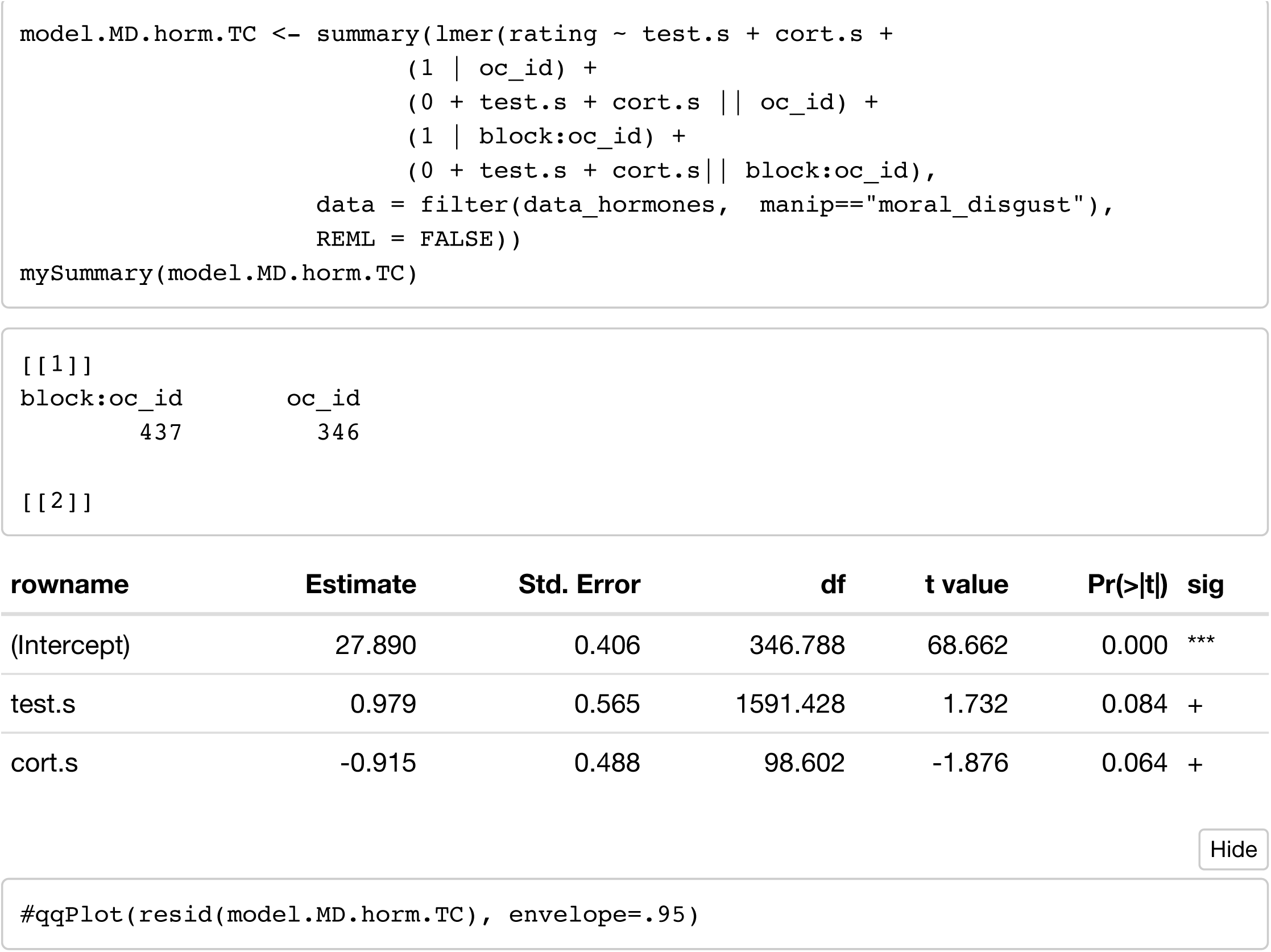

### Pathogen Disgust (controlling for session)

#### Model 1: pathogen_disgust ~ E + P + E x P

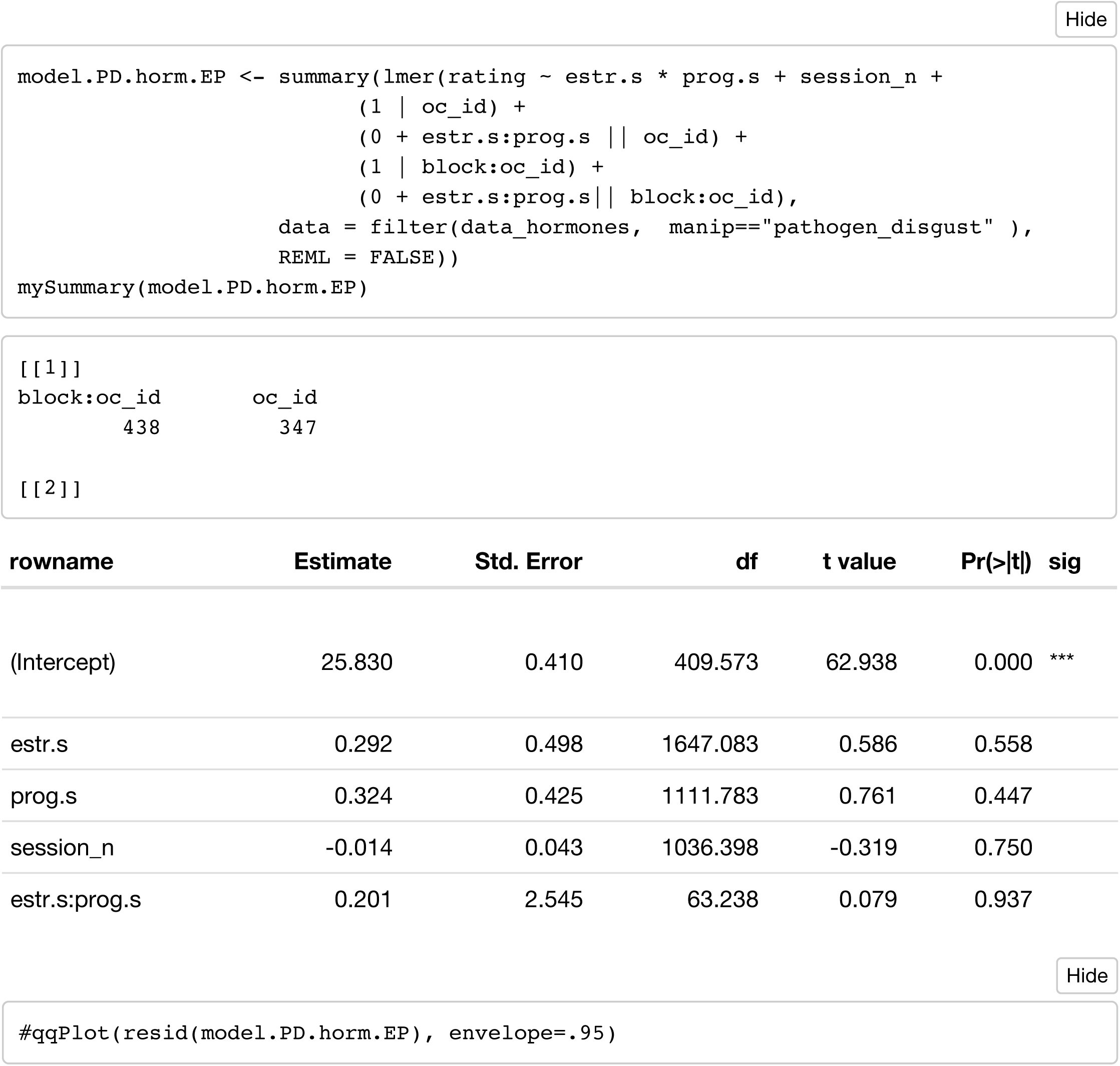

#### Model 2: pathogen_disgust ~ E + P + EP_ratio

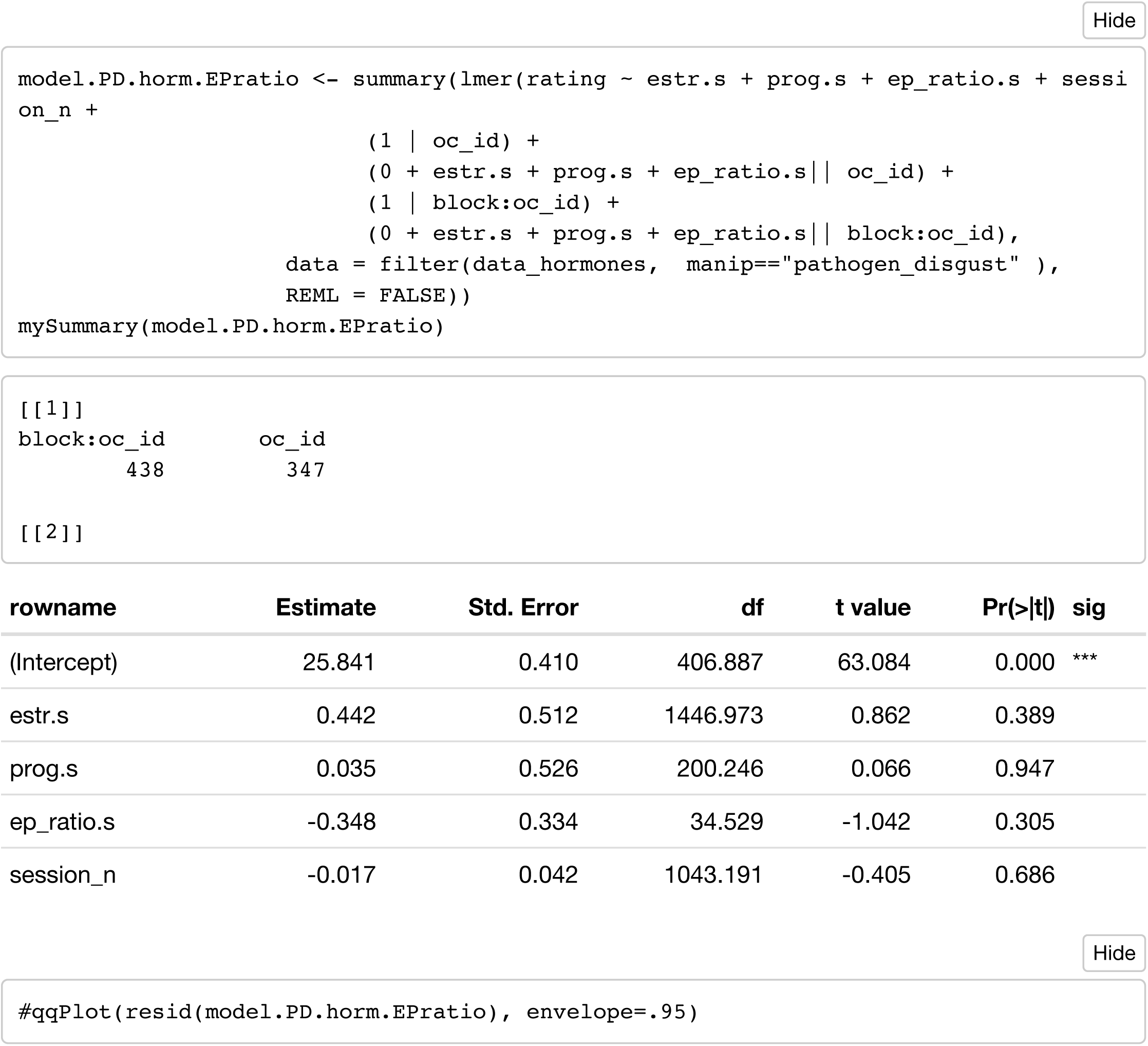

#### Model 3: pathogen_disgust ~ T + C

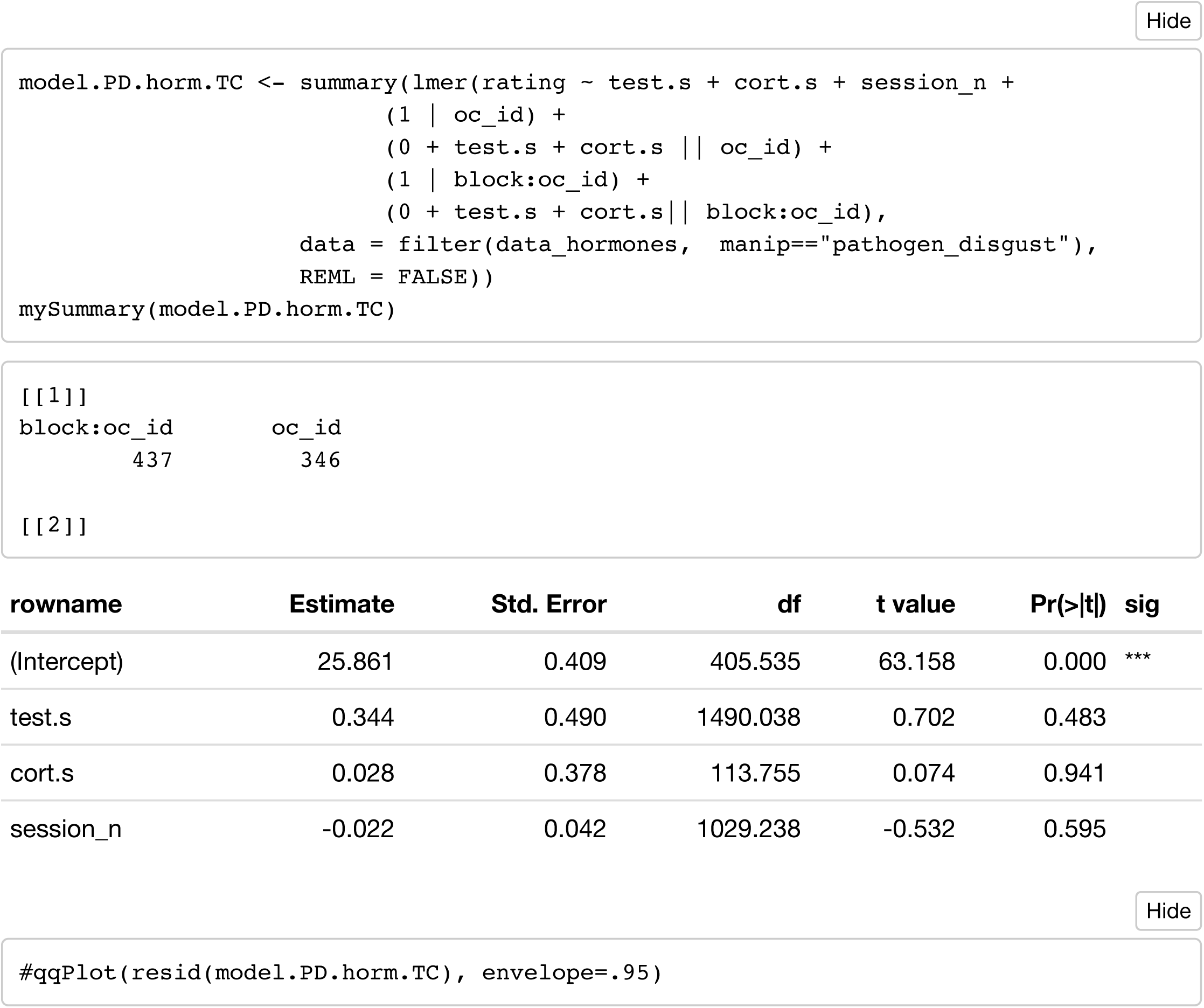

#### Model 4: pathogen_disgust ~ P

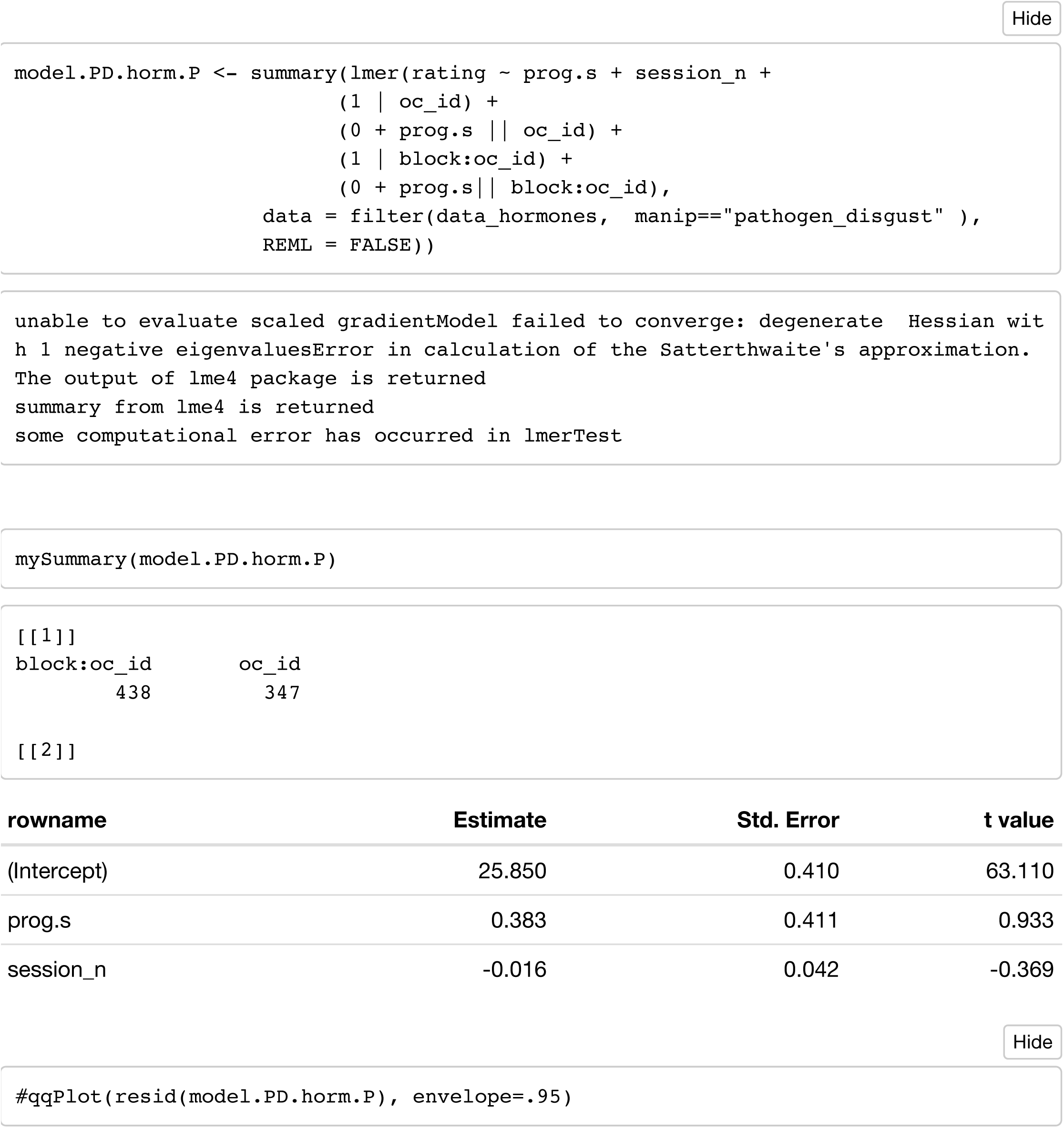

### Sexual Disgust (controlling for session)

#### Model 1: sexual_disgust ~ E + P + E x P

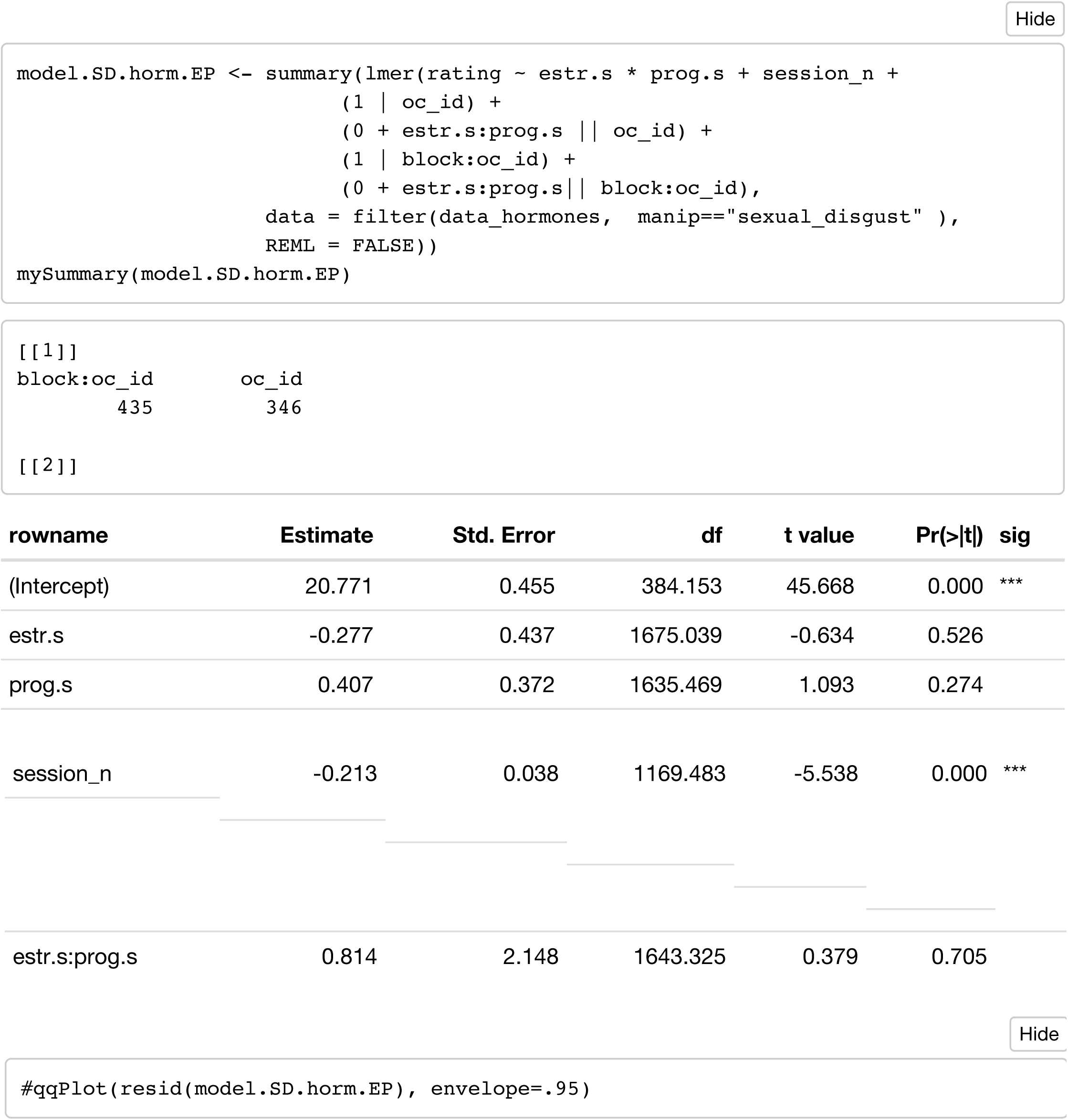

#### Model 2: sexual_disgust ~ E + P + EP_ratio

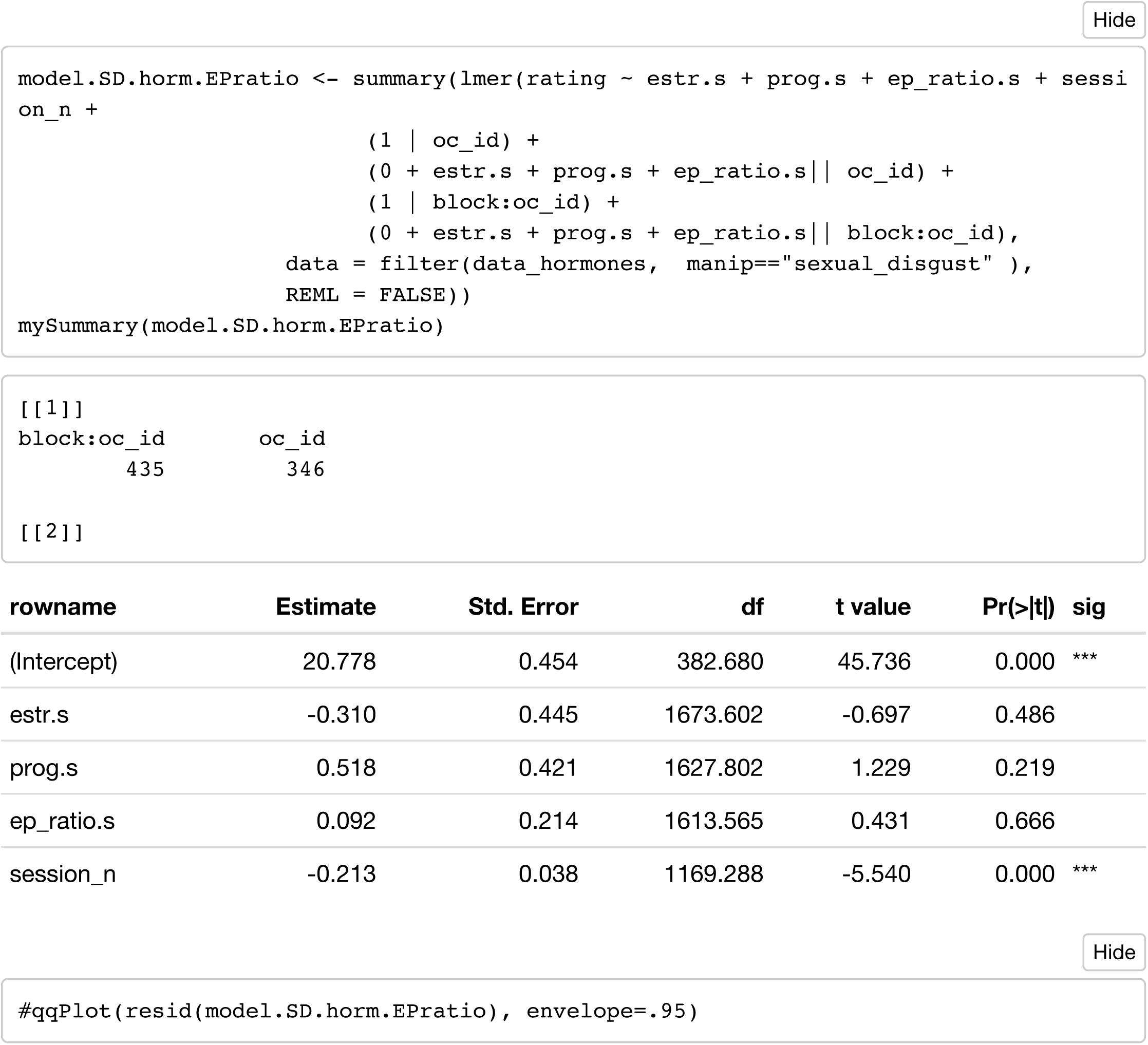

#### Model 3: sexual_disgust ~ T + C

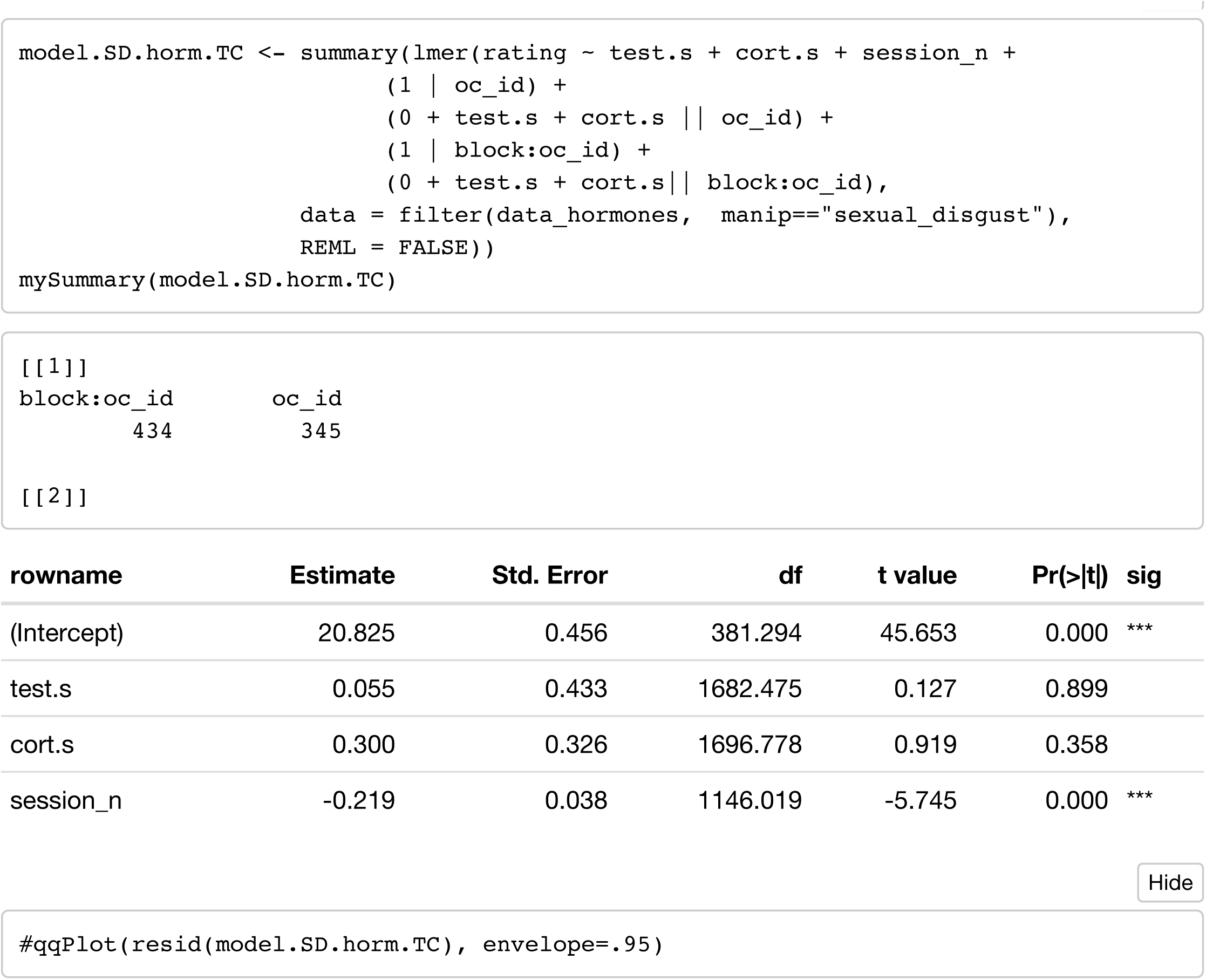

### Moral disgust (controlling for session)

#### Model 1: moral_disgust ~ E + P + E x P

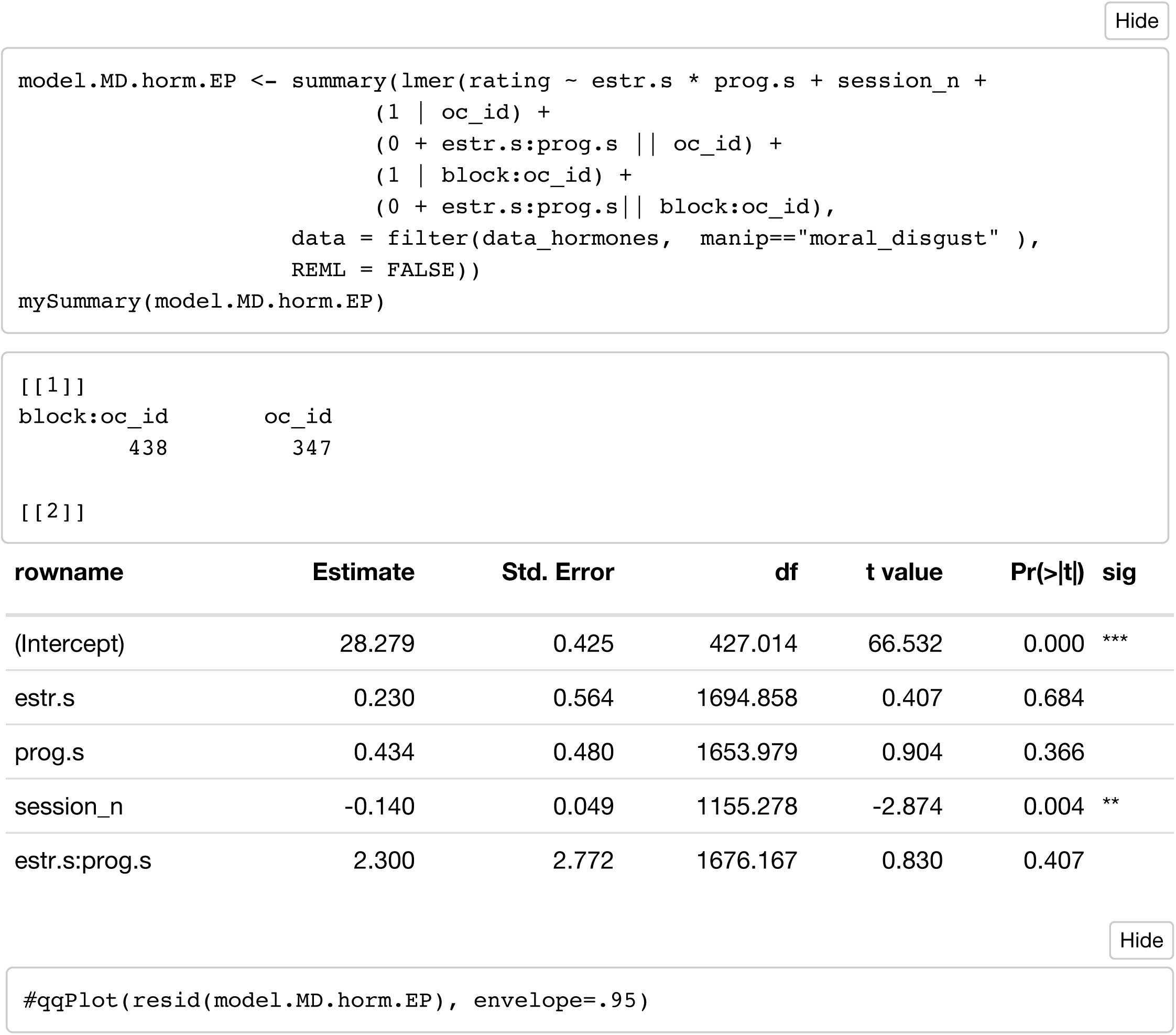

#### Model 2: moral_disgust ~ E + P + EP_ratio

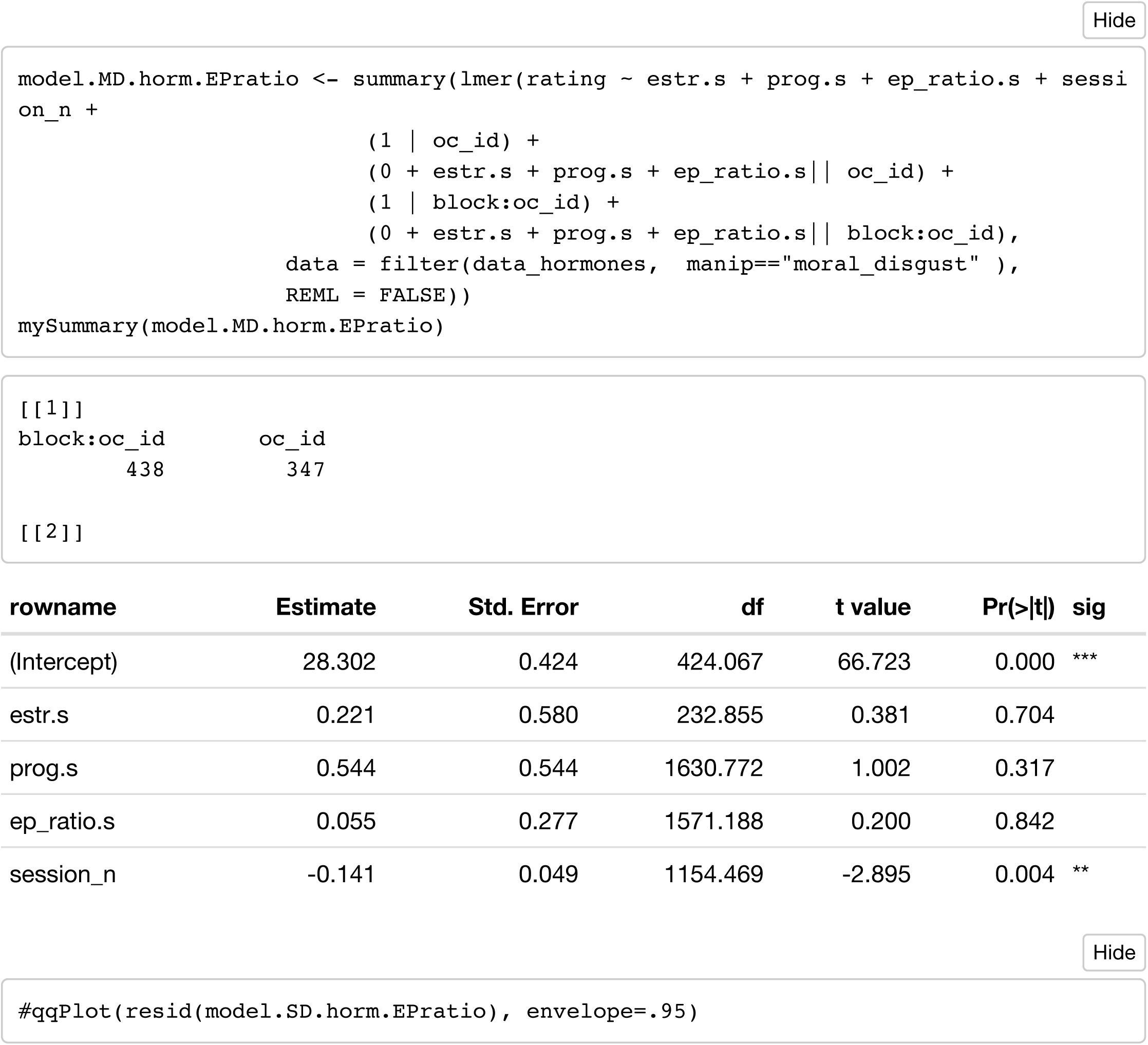

#### Model 3: moral_disgust ~ T + C

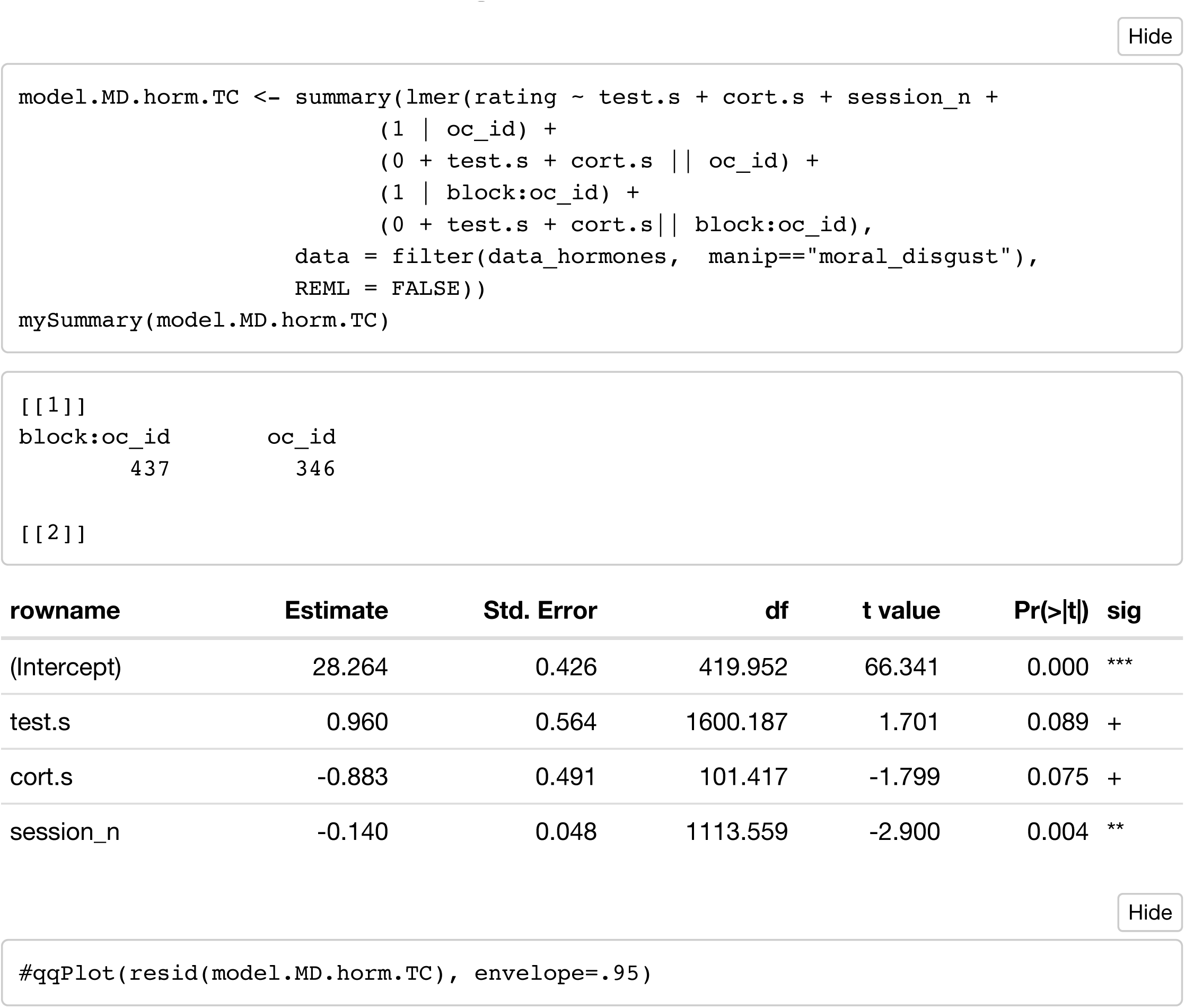

### Cross-sectional

#### Model 1: pathogen_disgust ~ P

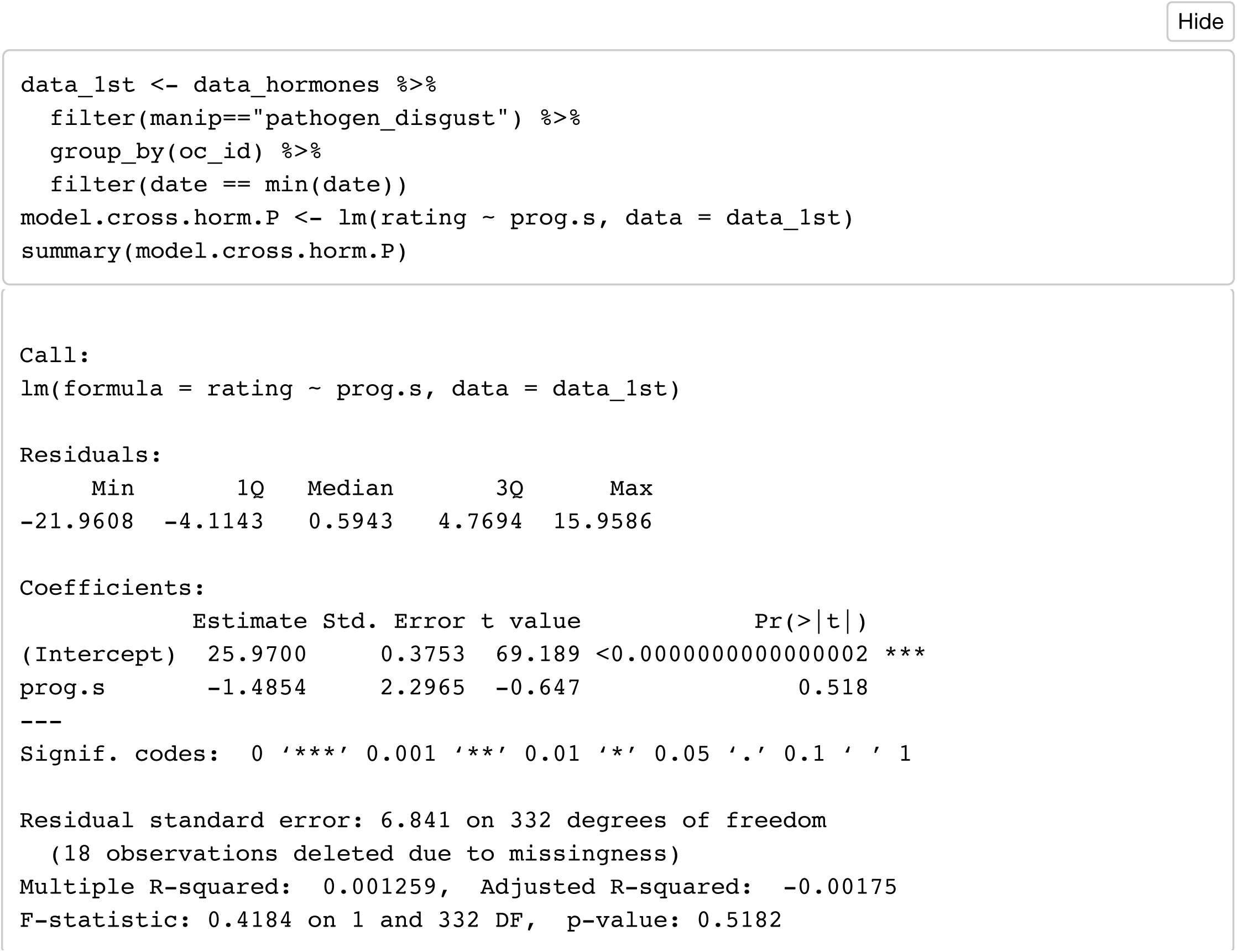

